# Integration of Morphometric and Machine Learning Approaches Strengthen Yield Prediction and Genetic Divergence Assessment in *Annona reticulata* under Semi-Arid Conditions

**DOI:** 10.64898/2026.05.15.725594

**Authors:** Vikas Yadav, Daya Shankar Mishra, Jagadish Rane, V.V. Apparao, Levitikos Dembure, Prakashbhai Ravat, Nezif Abajebal Abadura, Prabhat Kumar, Bismark Anokye, Anand Sahil, Poonam Devi, Peter Amoah

## Abstract

This study integrated morphometric characterization and machine-learning modelling to identify key predictors of yield in *Annona reticulata* under semi-arid conditions. Thirty-one canopy, fruit, seed, and biochemical traits were evaluated across 62 genotypes, revealing substantial phenotypic diversity, particularly in structural attributes such as tree growth nature and branch angle. Principal Component Analysis and hierarchical clustering differentiated genotypes into three ideotypes representing high-yielding, structurally stable, and quality-oriented groups. Random Forest modelling and SHapley Additive exPlanations (SHAP) interpretation consistently highlighted leaf breadth, leaf length, fruit shape, and pulp-associated traits as dominant yield predictors, underscoring the coordinated influence of source-sink balance. Integration of SHAP importances with trait stability (CV%) further revealed that moderately variable traits provide reliable selection indices. These findings demonstrate that yield performance is governed by multivariate trait networks rather than isolated descriptors. The proposed framework provides a robust basis for precision phenotyping and strategic parent selection to develop high-yielding, nutritionally enriched, and climate-resilient custard apple cultivars.

## Introduction

Fruit crops are well established as a key component of global horticulture, providing food and nutritional security, generating rural employment, and serving as a major source of raw materials for agro-industrial development [1,2]. Among them, *Annona reticulata*, also known as ramphal is a hardy perennial fruit tree native to the Indian subcontinent and parts of Southeast Asia. The species is uniquely adapted to arid and semi-arid environments, demonstrating resilience to drought, heat stress, and nutrient-poor soils, which positions it as a candidate for climate-resilient horticultural systems [3–5]. Its fruits, enriched with carbohydrates, minerals, and bioactive compounds, are valued for nutritional, medicinal, and economic purposes [6–8]. Therefore, the ecological adaptability and multifaceted utility of *Annona reticulata* highlight its potential as a strategically important fruit crop for sustainable production systems in the face of increasing climate variability.

Accurate yield prediction in perennial fruit crops is a complex but essential task that is increasingly being enhanced through the application of machine learning approaches. This yield prediction remains inherently complex due to a confluence of factors encompassing prolonged juvenile phases, irregular flowering and flowering time and sensitivity to environmental fluctuations [9,10]. These biological challenges render conventional statistical approaches inappropriate for underutilized crops such as ramphal. Fortunately, advances in artificial intelligence (AI), particularly machine learning (ML), offer promising avenues to address these challenges by enabling more accurate prediction, analysis, and optimization of complex agricultural systems. ML algorithms enable the modelling of high-dimensional, non-linear relationships among plant traits, environmental factors, and yield, thereby improving both the accuracy and interpretability of forecasting [11,12]. Beyond being a theoretical exercise, yield prediction provides practical value by supporting production planning, market forecasting, efficient resource allocation, and postharvest management, ultimately contributing to the profitability, sustainability, and long-term resilience of perennial fruit systems while enhancing farmer income [13,14].

Within this framework, morphometric traits, such as tree height, canopy spread, branching pattern, leaf and fruit morphology and pulp characteristics serve as reliable indicators of vegetative vigour and reproductive efficiency. Previous studies in mango, guava, grapes and some other tropical fruits have demonstrated strong correlations between these traits and yield outcomes, with attributes like canopy volume, stem girth, fruit shape and size acting as robust predictors [15–17]. When combined with weather variables such as temperature, rainfall, and relative humidity, morphometric descriptors can provide a holistic framework for yield modelling and prediction. Modern approaches such as regression analysis, principal component analysis (PCA), and advanced ML models such as Random Forests and neural networks have been demonstrated to enhance prediction accuracy by capturing complex interactions in many crops [18]. However, such integrative approaches remain largely unexplored in underutilized crops, highlighting a critical research gap that warrants further investigation. Against this backdrop, the present study was designed to (i) identify morphometric traits significantly associated with yield in ramphal, (ii) apply ML-based models to predict yield from trait datasets, and (iii) classify genotypes into performance groups using clustering approaches. The outcomes are expected to provide new mechanistic insight into how morphological traits collectively shape yield performance, enabling targeted breeding of compact, high-yielding ideotypes. These integrative results also strengthen the foundation for precision and climate-smart horticulture, supporting sustainable improvement and wider adoption of this underutilized perennial fruit crop.

## Material and methods

### Plant Material and Experimental Layout

The study utilized 23 ramphal genotypes maintained in the field gene repository at CHES, Panchmahal, Gujarat, India. It was conducted at the Central Horticultural Experiment Station (CHES), ICAR, Vejalpur, Panchmahal, Gujarat, India (22°41′N, 73°33′E; altitude 113 m above sea level) over two consecutive fruiting seasons (2023 and 2024). The experiment was laid out in a randomized complete block design (RCBD), with each genotype represented by 9–10-year-old trees established at a spacing of 5 × 5 m. The experimental site is characterized by a hot, semi-arid climate with an average annual rainfall of 630 mm. The soils of the experimental farm are predominantly shallow, with a clay-loam texture, a pH of approximately 6.7, and organic carbon content ranging from 0.43 to 0.47%. Standard recommended crop management practices were followed to ensure uniform growth and optimal tree performance [61,62]. To assess phenotypic diversity among the genotypes, 31 morphological traits were recorded (Fig. 1A and B) and systematically coded for analysis (Table 1).

**Fig. 1.**
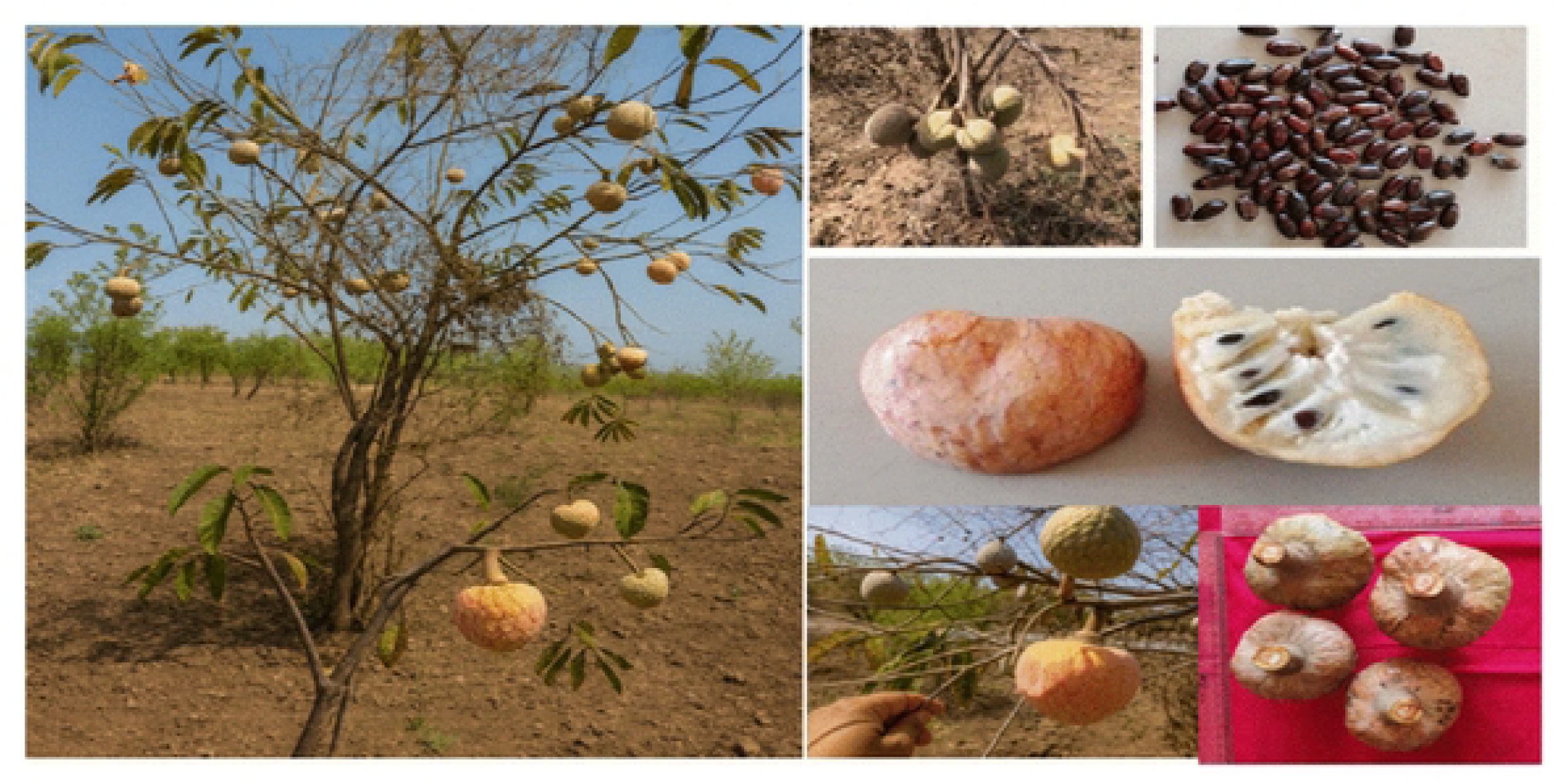
A: Morphological diversity of *Annona reticulata* genotypes showing representative tree, leaf, fruit, pulp, seed characteristics, and leaf variability among the studied accessions.

**Fig. 1B:**
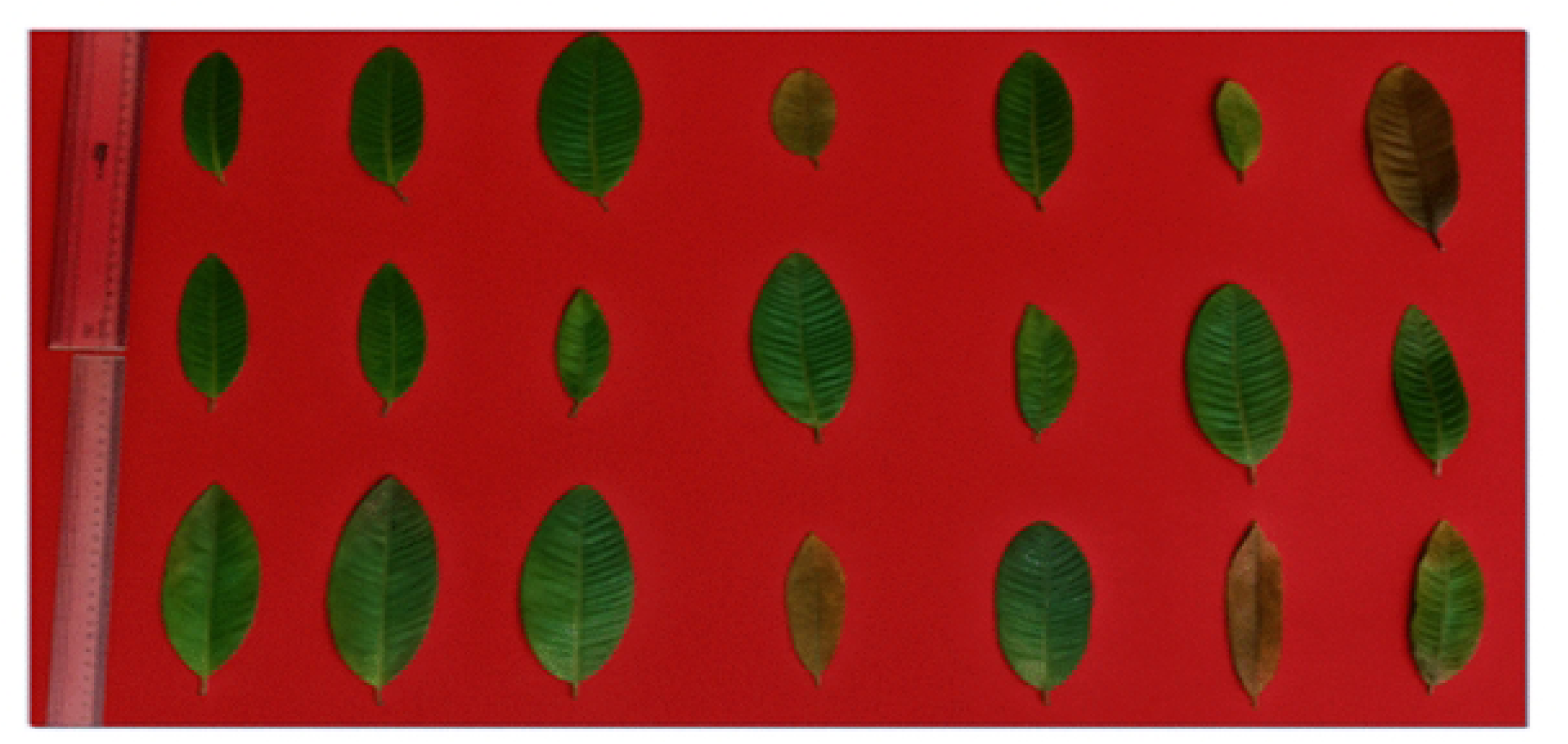
Variation in leaf morphology and phenotypic diversity among the evaluated *Annona reticulata* genotypes.

**Table 1:**
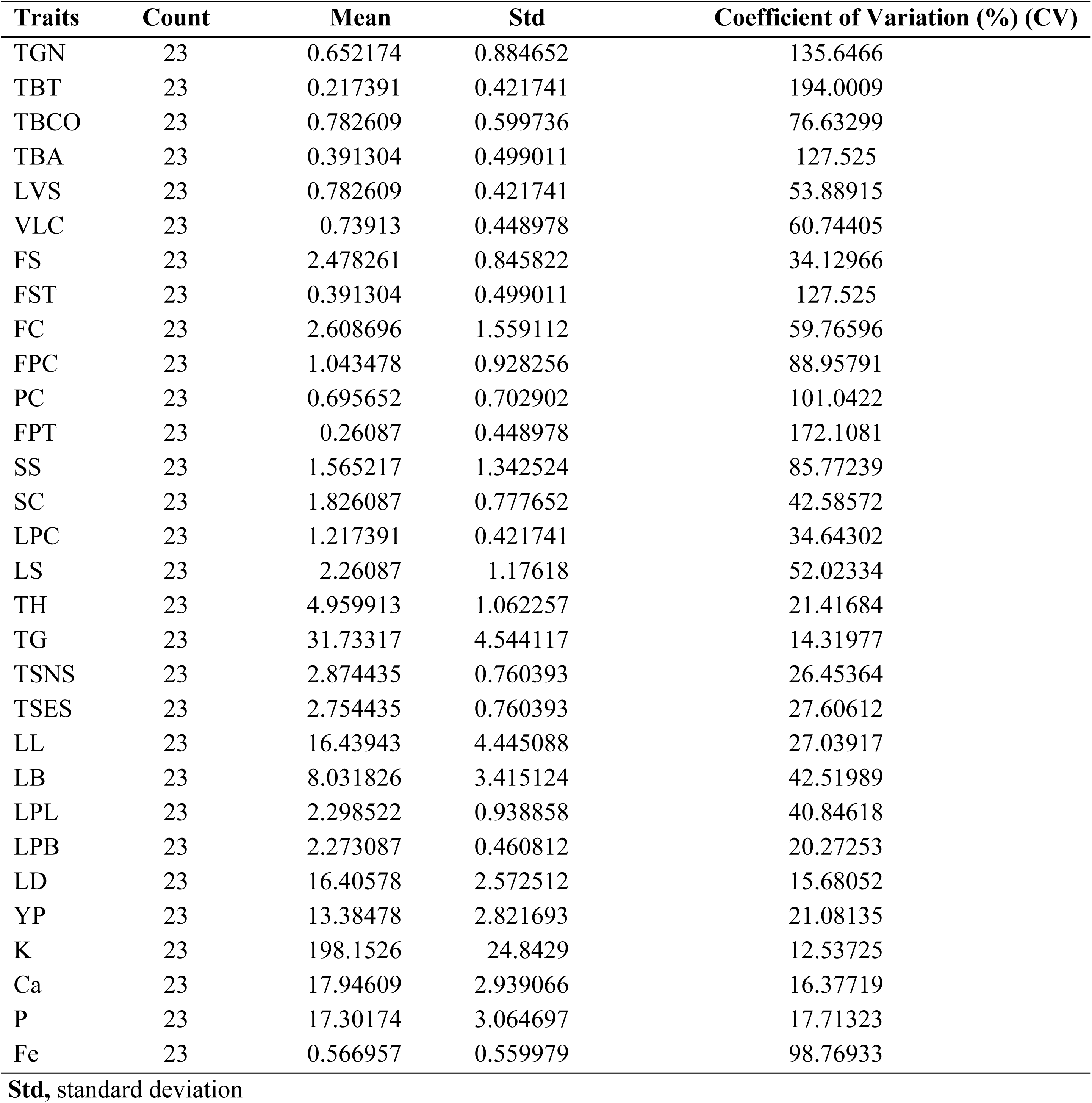
Descriptive statistics of morphological, yield, and biochemical traits of *Annona reticulata* genotypes used in this present study.

### Morphological Assessment of Tree, Leaf, and Fruit Traits

Morphological variation among the 23 ramphal genotypes was evaluated using 31 traits, covering vegetative, leaf, floral, fruit, and seed characteristics, including **TGN** – Tree Growth Nature, **TBT** – Tree Bark Texture, **TBCO** – Tree Bark Colour, **TBA** – Tree Branch Angle, **LVS** – Leaf Ventral Side, **VLC** – Vegetative Life Cycle, **FS** – Fruit Shape, **FST** – Fruit Surface Texture, **FC** – Fruit Colour, **FPC** – Fruit Pedicel Colour, **PC** – Pulp Colour, **FPT** – Fruit Pulp Texture, **SS** – Seed Shape, **SC** – Seed Colour, **LPC** – Leaf Petiole Colour, **LS** – Leaf Shape, **TH** – Tree height, **TG** – Tree girth, **TSNS** – Tree spread north-south, **TSES** – Tree spread east-west, **LL** – Leaf Length, **LB** – Leaf Breadth, **LPL** – Leaf Petiole Length, **LPB** – Leaf Petiole Breadth, **LD** – Leaf density, **YP** – Yield per Plant **K** – Potassium Content, **Ca** – Calcium Content, **P** – Phosphorus Content and **Fe** – Iron Content. Observations were recorded following standard descriptors developed for *Citrus* (IPGRI, 1999) and related species [20].

The tree traits (TGN, TG, TH, TBT, TBCO, TBA, TSNS, TSES and LVS) were measured before fruit harvest in the second week of July. Fully mature leaves (4-6 months old) from the middle portion of the current season’s growth were collected for leaf and leaflet observations (n = 50). and fruit traits were measured from fruits harvested between late November and Decemberr (n = 10). All samples were collected from 2-3 representative trees per genotype.

### Tree growth and leaf physical characters

Tree growth and yield characters, including tree height (TH), Tree girth (TG), and tree spread in North-South (SPNS) and East-West (SPES) directions were recorded following standard procedures. The TH and TG were measured before the fruit harvest in the second week of December. For leaf and leaflets observations, fully mature leaves from the middle portion of the current season’s growth, ensuring no signs of active growth were selected. Tree girth was used to calculate the SD using the formula: SD= π (d/2)2; where d= mean of E-W and N-S stock diameters [16]. Fruit yield per tree was determined by summing the fruits harvested during the fruiting season, from the last week of November to January.

### Statistical analyses

All data analyses were conducted using Python (version 3.9) with a suite of libraries tailored for statistical and machine learning tasks: pandas for data manipulation [23], NumPy for numerical computations [24], scikit-learn for machine learning and preprocessing [25], seaborn and matplotlib for visualization [26,27], and SHAP for model interpretability [28]. The data set comprising 186 observations of 62 wood apple genotypes (three replications each), included 32 morphological traits and yield (YT). Data were split into training (80%) and testing (20%) sets with a random seed of 42, and traits were standardized using Standard Scaler to normalize scales prior to analysis [25]. Five complementary analyses were performed to investigate trait-yield relationships. First, Pearson’s correlation coefficients were calculated and visualized as a half-triangle heatmap to assess linear associations between traits and yield [29]. Second, Principal Component Analysis (PCA) reduced the 32 traits to two components, with variance explained and trait loadings examined [30]. Third, a Random Forest Regressor (100 trees, random seed 42) modeled yield, with performance evaluated via Mean Squared Error and R², and feature importance extracted [31]. Fourth, SHAP values from the Random Forest were computed to interpret trait contributions to predictions [28]. Finally, K-Means clustering (k=3) grouped genotypes by traits, with yield distributions and cluster trait means analyzed [32].

## Result

### Frequency distribution of morphometric traits

The frequency distribution of qualitative traits among 23 *Annona reticulata* (ramphal) accessions revealed considerable morphological diversity across genotypes (Fig 2). The majority of trees exhibited an upright growth habit (14), followed by spreading (6) and semi-spreading (3). Smooth cylindrical bark was predominant (18), while rough bark was less frequent (4), suggesting less genetic diversity was observed in trunk surface characteristics. Like Whitish-grey bark colour (14 accessions) and coffee-coloured **(7)** or light-brown **(2)**, implying moderate pigmentation variability. Branch orientation varied moderately from narrow branch angles (14) to wide angles (9), reflecting differences in canopy openness and light interception potential might be influencing yield. The leaf ventral surface was predominantly dark green glabrous (18), and all accessions exhibited a light green, hair-spreading dorsal side (23). Most genotypes were semi-deciduous (17), with only six deciduous types, indicating seasonal adaptability and sustained life during mild stress periods. In fruit morphology, much variability was observed like spherical fruits (10), oblong (8), heart-shaped (3), and irregular (2) forms. Smooth fruit surfaces (14) were more prevalent than coarse textures (9), while fruit colour ranged such as pinkish-yellow (7), light brownish (6), pale yellowish (5), yellowish (3), and yellowish-red (2) showing broad pigment diversity and market-oriented variation. Pedicel colour also varied, with whitish-grey (10), grayish (9) and brownish (4). The pulp colour variability distribution dominance of pinkish-creamy (10) and creamy (10) accessions, followed by a few pinkish-yellow (3) types. Seed morphology exhibited broad variation, with oblong (7) and ovoid (6) shapes being most frequent, alongside spheroid (4), cuneiform (3), spheroid (2), and oval (1) forms. Although in cloure less diversity observed like Dark brown seeds (10), brown (9) and blackish-brown (4) suggesting modest pigment-based genetic diversity. Leaf attributes were noted relatively uniform like petiole colour was primarily dark green (18) and light green (5). Leaf shape showed balanced representation of oblong (7), obovate (6), lanceolate (6), and ovate (4). Overall, much genetic morphological diversity was observed, particularly in fruit, leaf and seed characteristics.

**Figure 2:**
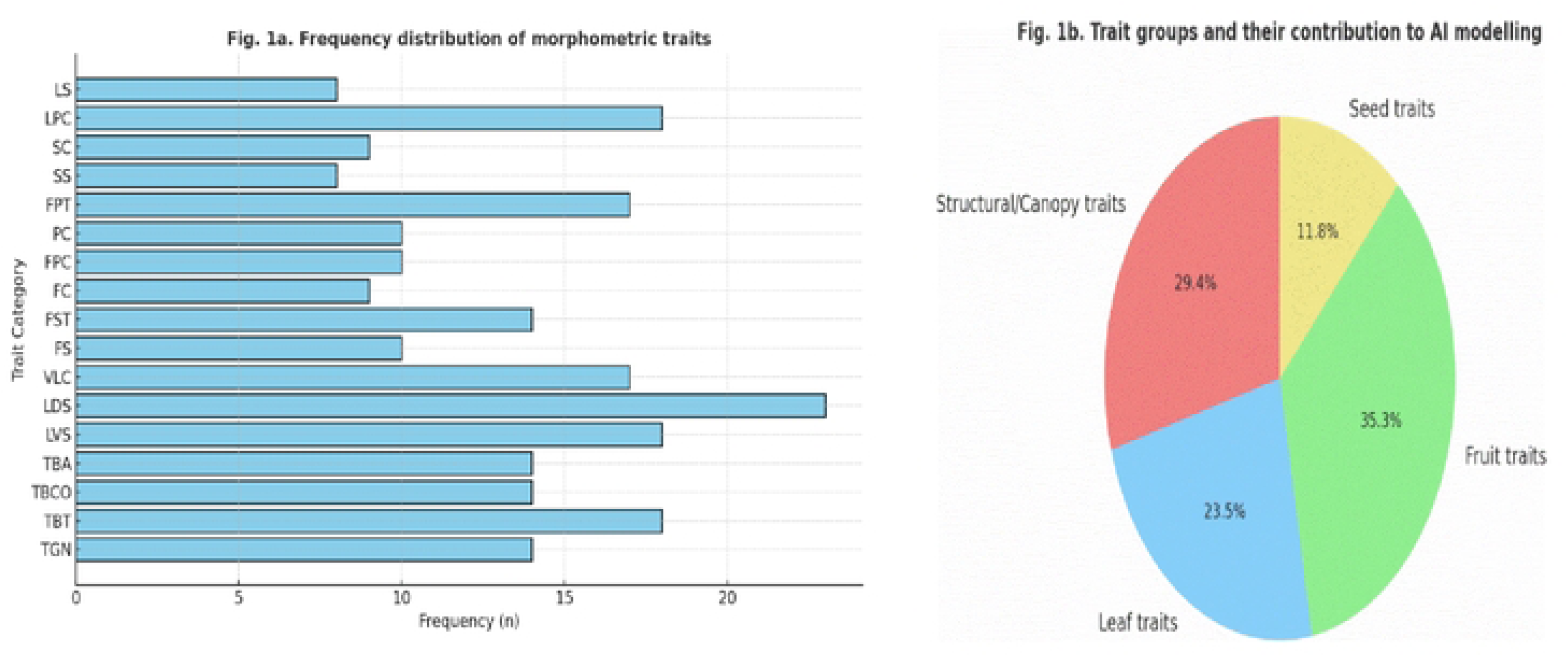
Distribution patterns of morphometric traits and their relative contribution to artificial intelligence-based predictive modelling among 23 ramphal genotypes.

### Descriptive statistics analysis

The coefficient of variation (CV%) analysis revealed substantial phenotypic diversity among the measured morphological, fruit, seed, and biochemical traits in *Annona reticulate* (Fig. 3), indicating a broad genetic base and strong scope for selection. The highest variability was observed in TBT (194.0%), TGN (135.6%), and FST (127.5%). These traits reflect pronounced architectural divergence among genotypes, likely driven by environmental adaptation and differential growth vigor under semi-arid conditions. Such traits may serve as promising selection criteria for identifying ideotypes with desirable canopy architecture and superior fruit textural quality. In contrast, comparatively lower variability was recorded for fruit and seed morphological traits, including Seed Shape (SS; 85.8%), Fruit Pedicel Colour (FPC; 89.0%), LS (52.0%), LB (42.5%), LP (40.8%), and SC (42.6%), indicating relatively greater phenotypic stability among the evaluated genotypes. This pattern indicates that reproductive traits possess sufficient diversity to support the improvement of consumer-preferred characteristics, while also offering opportunities for selecting genotypes with efficient canopy architecture and enhanced photosynthetic performance. In contrast, biochemical traits such as K (12.5%), Ca (16.4%), P (17.7%), and Fe (98.8%) exhibited comparatively lower CV values, suggesting stronger genetic control and relatively reduced environmental influence. Yield per plant (YP; 21.1%) also showed considerable variability, reflecting substantial diversity in yield performance among the evaluated accessions.

**Figure 3:**
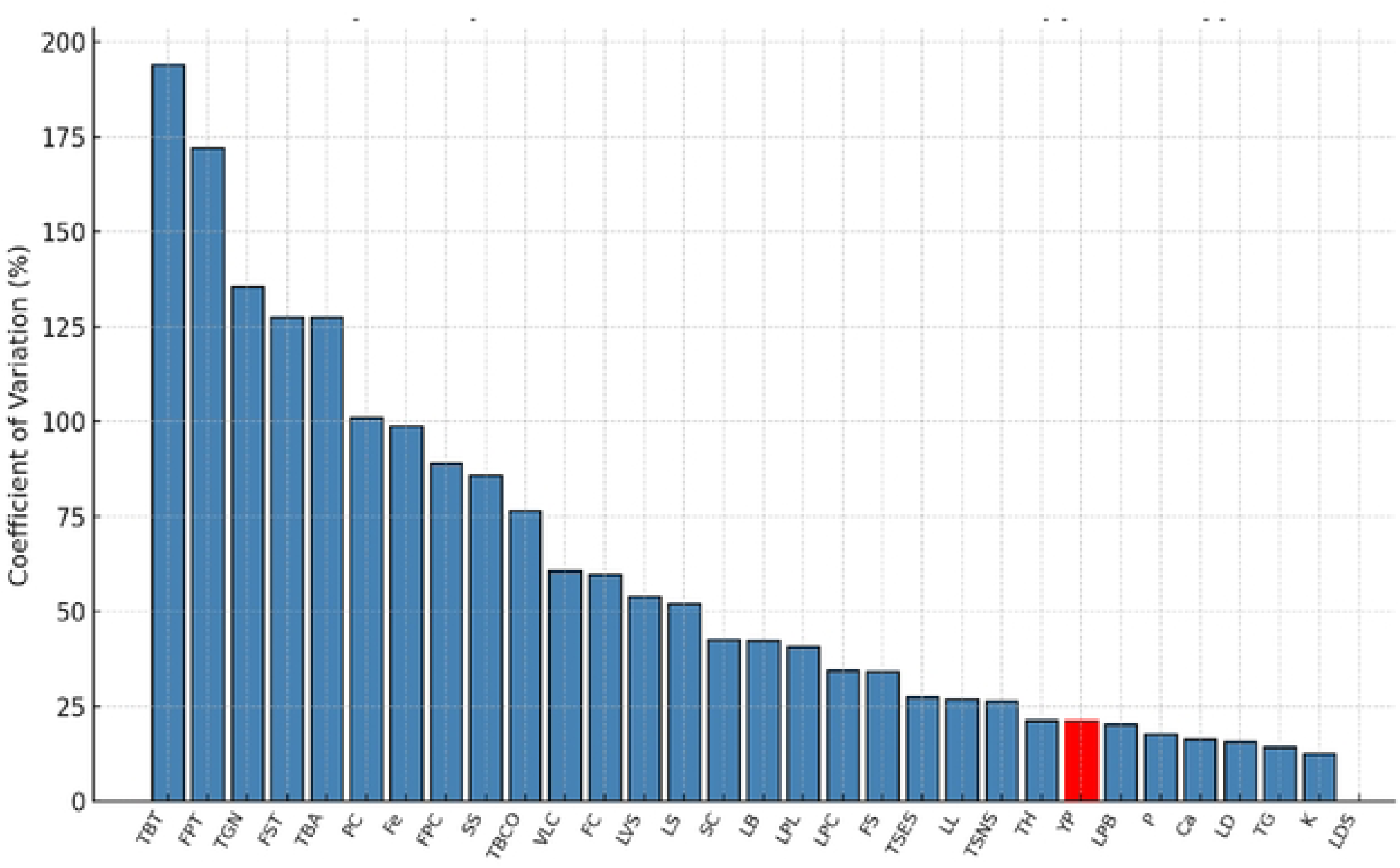
Trait-wise phenotypic variability expressed as coefficient of variation (CV%) for morphometric, fruit, seed, biochemical, and yield traits among 23 *Annona reticulata* genotypes.

The variability patterns observed through CV% align closely with the PCA and SHAP-based predictions, reinforcing the biological relevance of key determinants of yield. Traits exhibiting high CV (LB), (LL), (FS), and petiole dimensions also showed impact yield predictors in SHAP analysis, suggesting that stable yet sufficiently variable traits best explain yield variation. PCA further supported this relationship, with these traits showing high positive loadings on PC1 and PC2, the components most associated with canopy vigor, fruit morphology, and yield performance. Although traits such as TBT, FPT, and TGN contributed minimally to SHAP importance and PCA variance, they exhibited limited predictive reliability and relatively weak influence on yield prediction performance. Together, the CV–PCA–SHAP convergence confirms that moderately variable morphological traits with strong functional roles in source–sink balance are the most meaningful predictors for selecting high-yielding genotypes.

### Correlation analysis

The correlation matrix (**Fig4 and Table 2**) revealed clearer and more biologically meaningful relationships among morphological and yield traits of *Annona reticulata*. Strong positive correlations were observed between TGN and TBA (*r* = 0.94), indicating that spreading tree forms tend to exhibit wider branch orientation. Similarly, a positive correlation was also observed between LVS and Ca (*r* = 0.41*), implying efficient nutrient assimilation in structurally robust leaves. Conversely, TGN was negatively correlated with Tree Bark Colour (TBCO) (*r* = – 0.66*), showing that upright trees often display darker bark colour. Fruit-related traits showed moderate correlations, particularly between FST and PC (*r* = 0.42*), and between FC and YP (*r* = 0.30*) indicating that good shape fruits may also possess higher productivity potential. Nutrient traits like K, Ca, P, and Fe were positively correlated (*r* = 0.28–0.41*), reflecting synergistic nutrient accumulation patterns.

**Figure 4:**
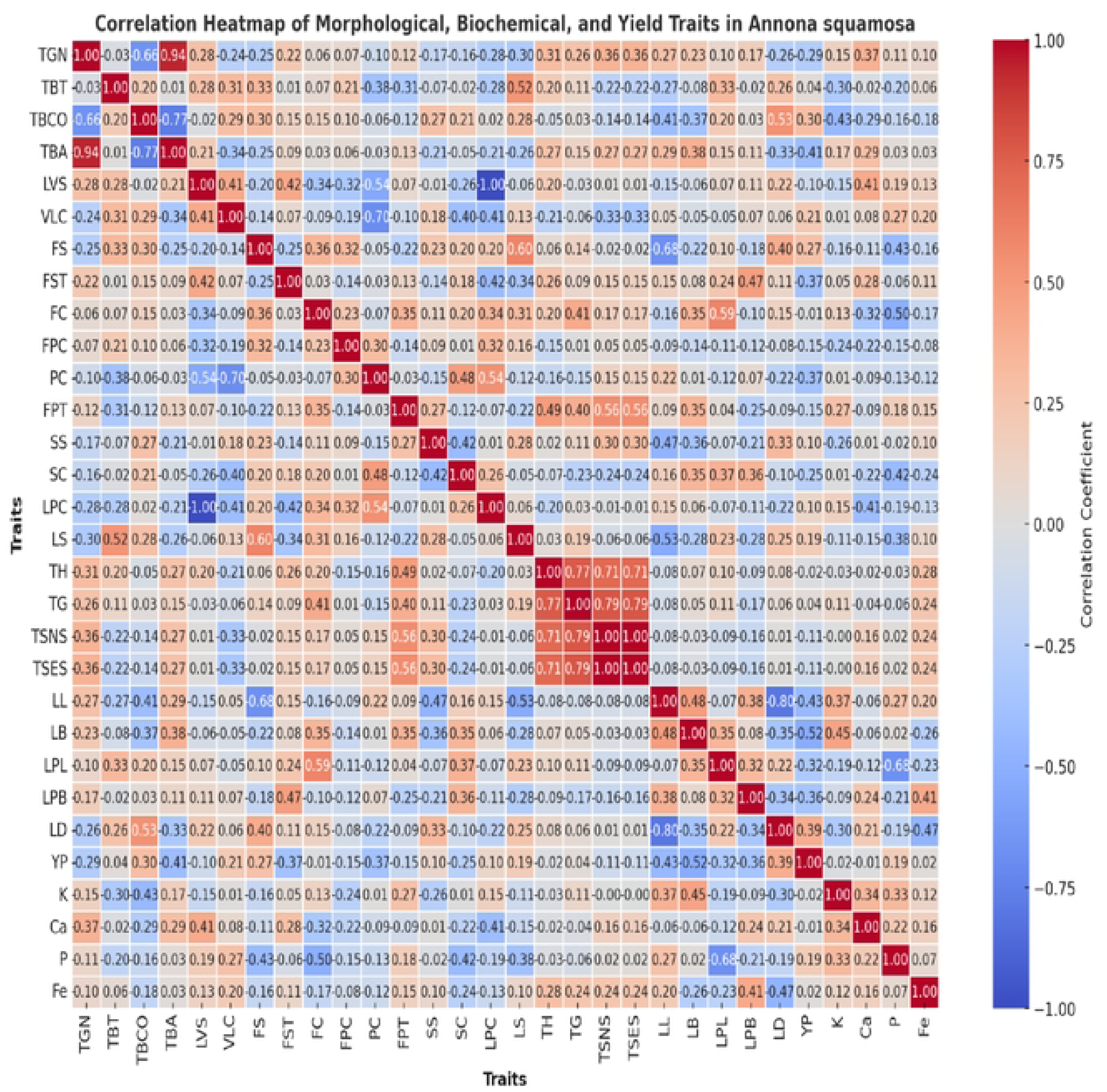
Correlation heatmap illustrating the relationships among morphological, yield, and biochemical traits of *Annona reticulata* genotype.

**Table 2:**
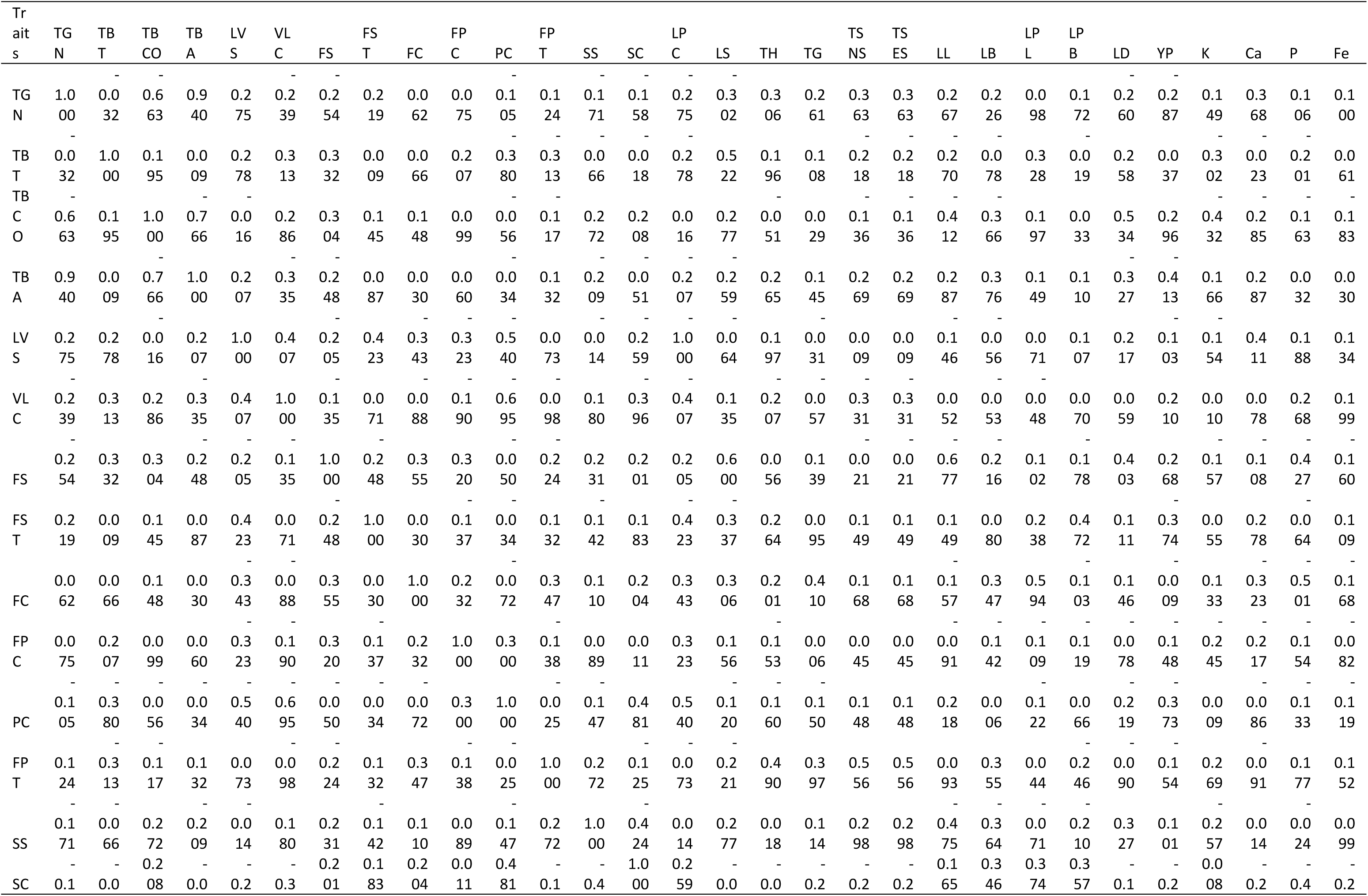
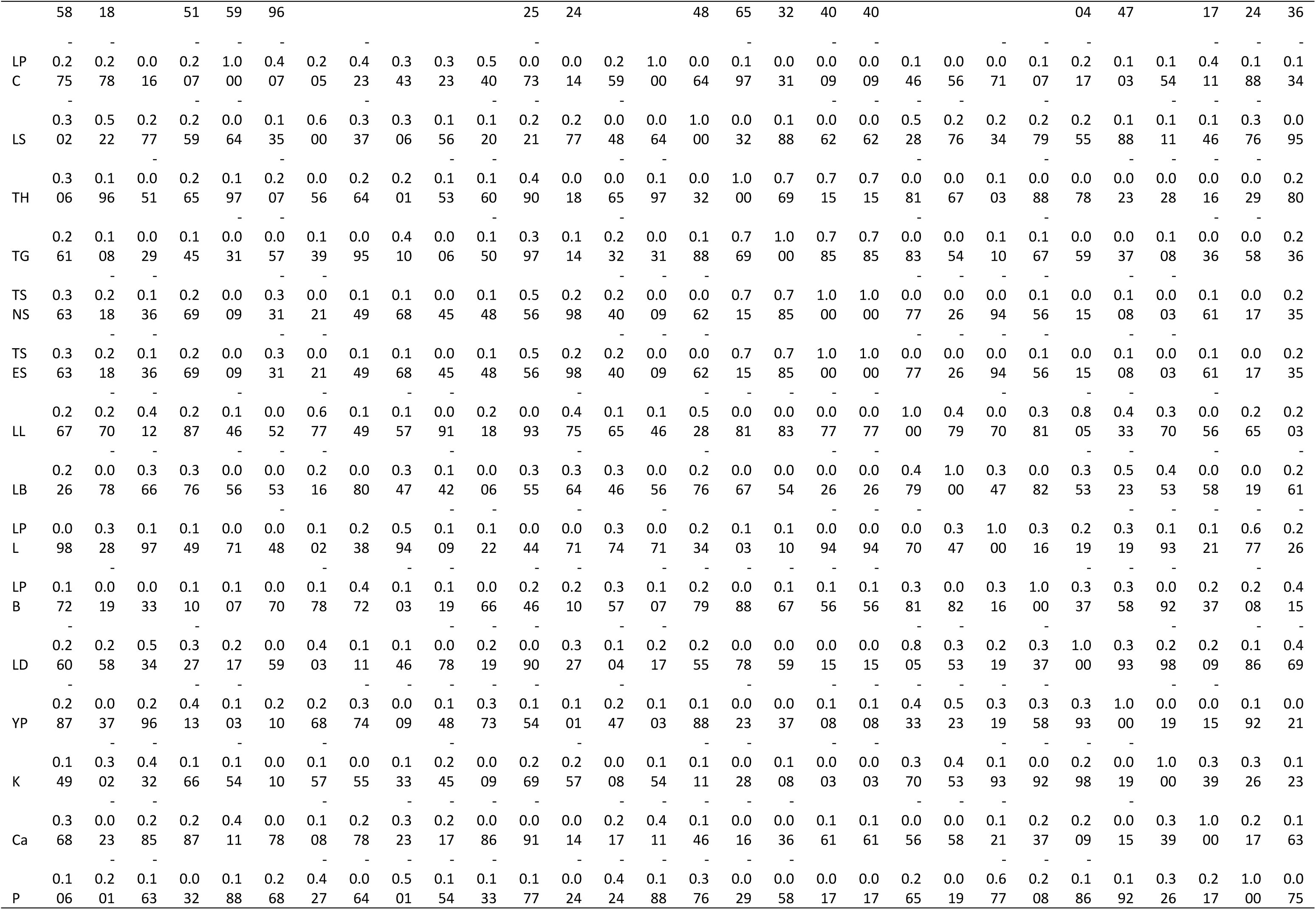
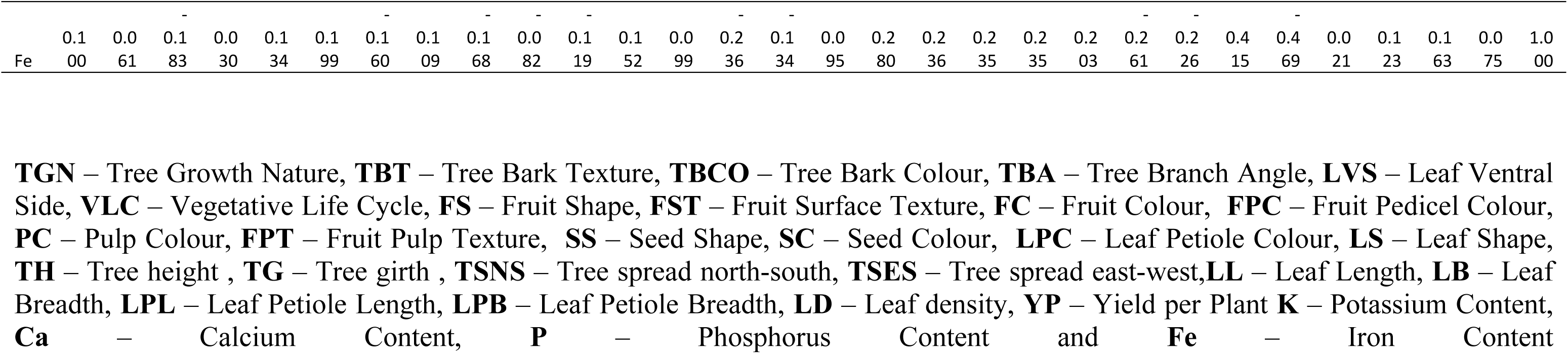
Correlation Matrix for morphological, yield, and biochemical traits in *Annona reticulata* genotypes.

### Principal Component Analysis (PCA)

Principal Component Analysis (PCA) was performed to determine the key morphological and yield-related traits contributing to genetic variability and those morphological traits much influencing yield traits among *Annona reticulata* genotypes (**Fig. 5 and Table 3,4,5**). The first two principal components (PC1 and PC2) together explained 32.5% of the total variance, with PC1 accounting for 17.9% and PC2 contributing 14.6%, indicating that these components effectively captured the major patterns of phenotypic diversity among the genotypes. PC1 was largely govern by tree architecture and fruit morphology, showing high positive loadings for traits such as *TGN*, *TBA*, and *FST*. This axis separated genotypes with spreading canopies, wide branch angles, and smoother fruit surfaces toward the positive direction. PC2, on the other hand, represented seed and yield-related attributes, with notable contributions from *SW*, and *YP*. The two-dimensional ordination clearly differentiated genotypes based on productivity potential. Genotypes plotted toward the upper right quadrant (positive PC1 and PC2) exhibited superior morphological balance and higher yield. In contrast, those positioned along the negative side of PC2 represented genotypes with compact canopy structure, smaller fruits, and comparatively lower yield performance. The color gradient in the PCA plot reflects Yield per Plant (YP), where warmer (yellow-orange) hues correspond to higher productivity and cooler (purple-blue) hues to lower yields. The top-yielding genotypes (Top-1, Top-2, Top-3) clustered closely within the positive region of both PC1 and PC2, confirming their superior performance across multiple trait dimensions. However, low-yielding genotypes (Low-1, Low-2, Low-3) appeared in the lower-left quadrant, suggesting less favorable combinations of fruit and canopy traits. The PCA score matrix revealed that high-yielding genotypes exhibited elevated positive scores for both PC1 and PC2, indicating that trait combinations such as wider branch angles, higher yield, smoother fruit surface texture, and related morphological characteristics collectively contributed to enhanced productivity. While, low-scoring genotypes along PC2 were characterized by narrow branching and pulp colour, yield, emphasizing the physiological trade-off between vegetative compactness and fruit yield.

**Figure 5:**
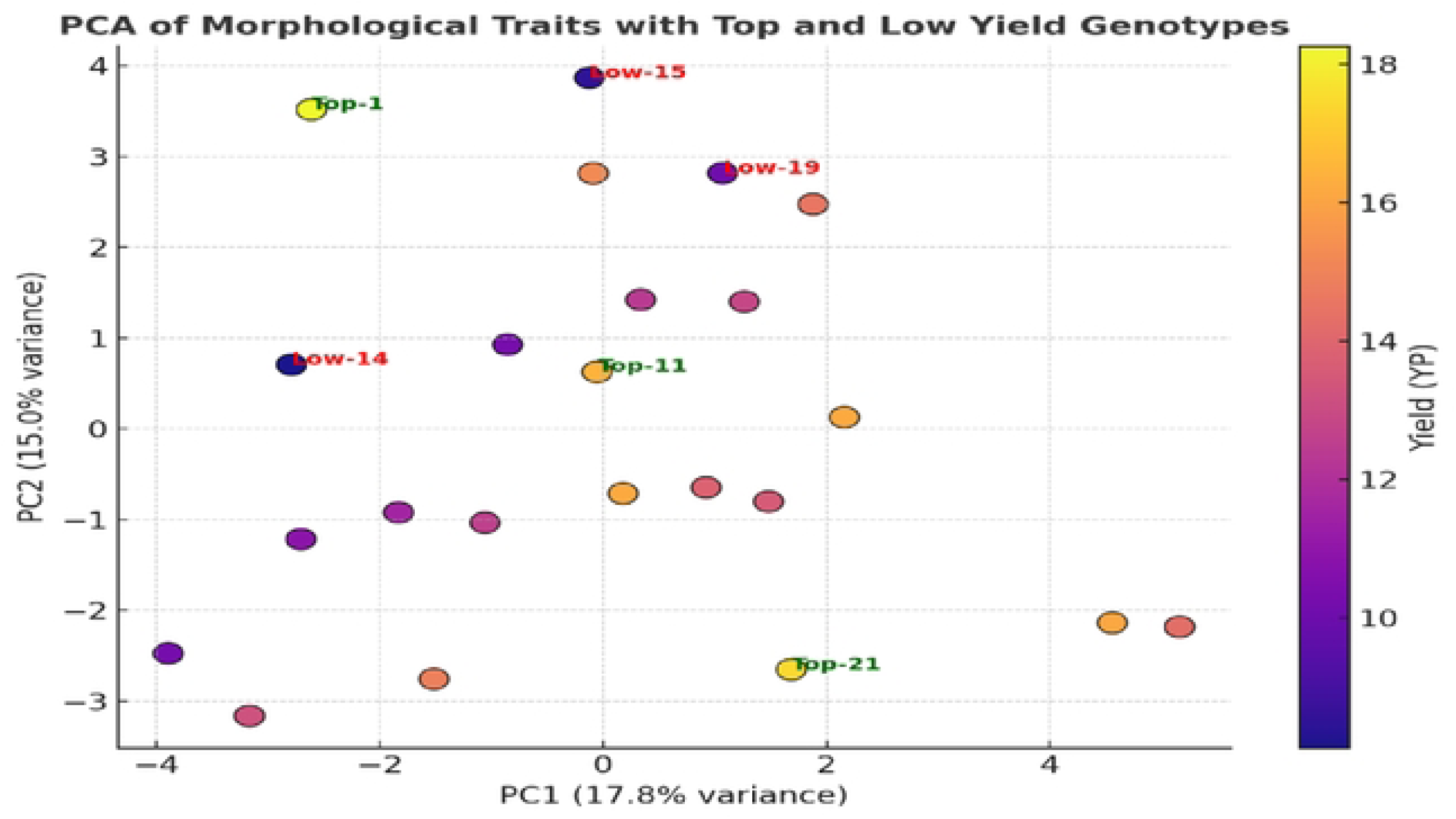
Principal Component Analysis plot showing distribution of *Annona reticulata* genotypes based on morphological and yield traits. The coloured gradient represents yield per plant (YP). Bold circular markers indicate genotypes; high-yielding accessions (Top-1-Top-3) are labeled in green and low-yielding ones (Low-1-Low-3) in red. PCI and PC2 explain 17.9 % and 14.6 % of total variance, respectively.

**Table 3:**
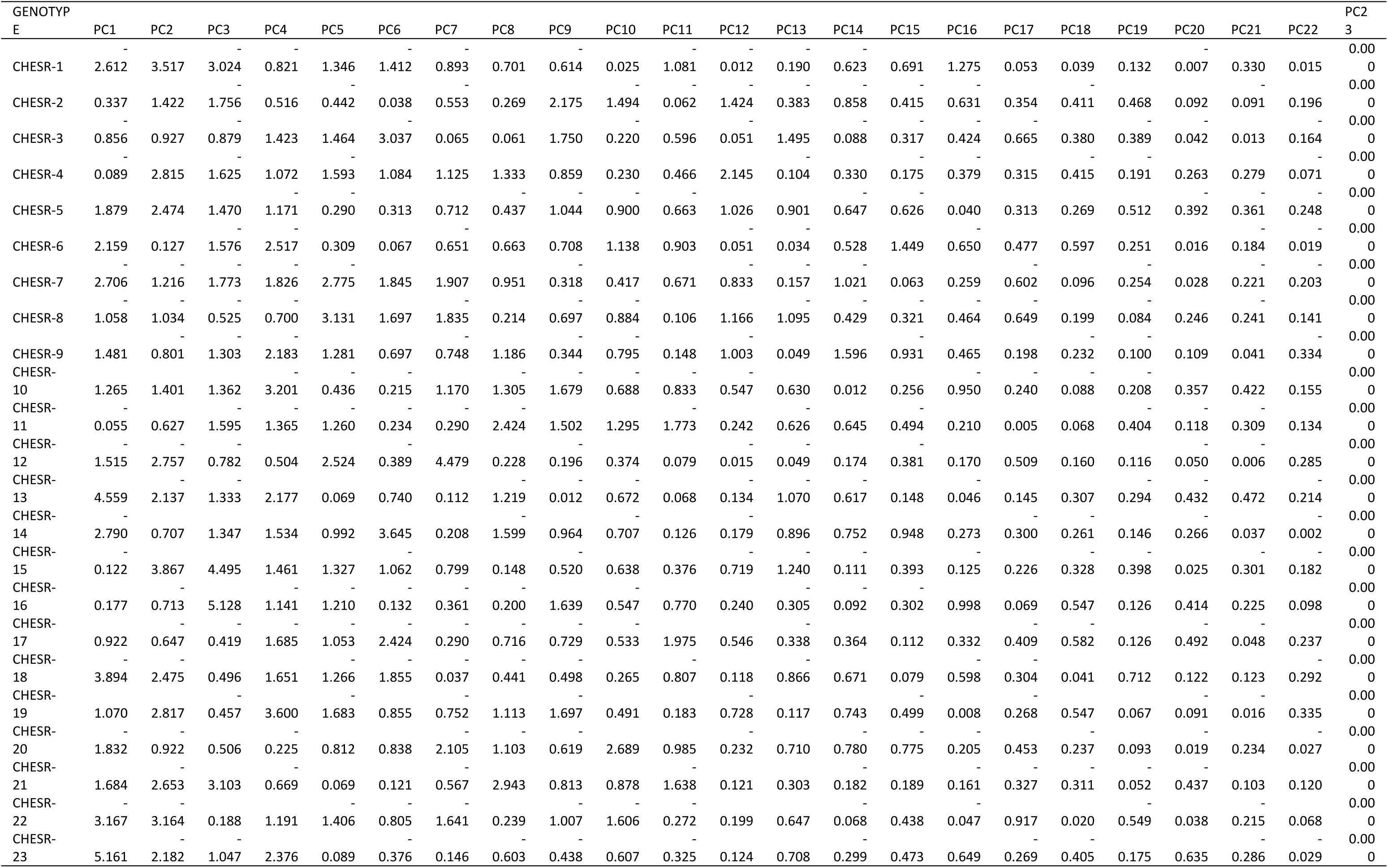
PCA component scores of *Annona reticulate* genotypes showing the contribution of principal components and yield performance.

**Table 4:** Correlation coefficients between principal component scores and yield per plant (YP) in *Annona reticulata* genotypes.

| Component | Correlation |
| --- | --- |
| YP | 1 |
| PC1 | 0.381805 |
| PC12 | 0.35217 |
| PC16 | 0.333778 |
| PC17 | 0.286025 |
| PC21 | 0.224937 |
| PC8 | 0.175695 |
| PC22 | 0.172936 |
| PC9 | 0.166391 |
| PC18 | 0.127472 |
| PC15 | 0.061149 |
| PC10 | 0.053745 |
| PC19 | 0.011735 |
| PC5 | -0.02255 |
| PC7 | -0.02938 |
| PC20 | -0.05914 |
| PC14 | -0.11181 |
| PC13 | -0.11933 |
| PC6 | -0.15165 |
| PC2 | -0.15699 |
| PC3 | -0.20273 |
| PC11 | -0.31655 |
| PC4 | -0.39313 |
| PC23 | 0 |

**Table 5:** Eigenvalues, explained variance, and cumulative variance of principal components derived from morphometric, yield, and biochemical traits in *Annona reticulata* genotypes.

| Principal Component | Eigenvalue | Explained Variance (%) | Cumulative Variance (%) |
| --- | --- | --- | --- |
| PC1 | 2.091117 | 8.696516 | 8.696516 |
| PC2 | 1.047178 | 4.354992 | 13.05151 |
| PC3 | 1.046713 | 4.353059 | 17.40457 |
| PC4 | 1.046683 | 4.352936 | 21.7575 |
| PC5 | 1.046445 | 4.351945 | 26.10945 |
| PC6 | 1.045971 | 4.349976 | 30.45942 |
| PC7 | 1.045906 | 4.349703 | 34.80913 |
| PC8 | 1.045787 | 4.349207 | 39.15833 |
| PC9 | 1.045742 | 4.349022 | 43.50736 |
| PC10 | 1.045675 | 4.348745 | 47.8561 |
| PC11 | 1.045601 | 4.348434 | 52.20454 |
| PC12 | 1.045481 | 4.347935 | 56.55247 |
| PC13 | 1.045387 | 4.347545 | 60.90002 |
| PC14 | 1.045275 | 4.34708 | 65.2471 |
| PC15 | 1.045221 | 4.346857 | 69.59395 |
| PC16 | 1.045047 | 4.346133 | 73.94009 |
| PC17 | 1.04493 | 4.345644 | 78.28573 |
| PC18 | 1.044855 | 4.345332 | 82.63106 |
| PC19 | 1.044473 | 4.343743 | 86.97481 |
| PC20 | 1.044247 | 4.342804 | 91.31761 |
| PC21 | 1.044053 | 4.341996 | 95.6596 |
| PC22 | 1.043668 | 4.340396 | 100 |

### Clustering analyses and yield distribution of clusters

Cluster analysis integrating morphological, fruit, seed, and biochemical traits revealed three well-defined phenotypic groups with distinct productivity profiles among *Annona reticulata* genotypes (**Table 6 and Fig 6,7,8**). Cluster 1 emerged as the superior group, combining semi-spreading canopies (TGN = 1.74), smoother fruit surfaces (FST = 0.86), desirable fruit shape (FS = 2.98), and elevated mineral levels (Ca = 1.86; P = 1.47 mg 100 g⁻¹). These attributes translated into the highest yield, as confirmed by the boxplot distribution, with a median of 16.3kg plant⁻¹, a narrow IQR of 3.7 kg, and upper whiskers reaching 17.3–18.4 kg plant⁻¹. Such uniform performance indicates efficient resource use and strong genetic consistency, making Cluster 1 ideal for selecting high-yielding, stable parental lines. Cluster 0 displayed a moderate mean yield of 13.19 kg plant⁻¹and a median of 14.2 kg plant⁻¹, accompanied by a wider IQR (5.1 kg; 10.1–18.3 kg plant⁻¹). Higher mean Tree Bark Colour (1.52), Tree Branch Angle (0.91), and Seed Shape (2.36), along with above-average TSS (25.6 °Brix), indicate a group with balanced canopy structure and favorable organoleptic characteristics. The broader yield range reflects greater genetic heterogeneity, offering a diverse pool of adaptable and stress-tolerant genotypes suitable for semi-arid cultivation. Cluster 2, although exhibiting enriched fruit and seed quality attributes--highest pulp colour (PC = 1.88), seed colour (SC = 2.97), and iron content (Fe = 3.26 mg 100 g⁻¹)--recorded the lowest yield with a mean of 11.05 kg plant⁻¹ and median of 11.6 kg plant⁻¹, accompanied by a compact IQR of 3.0 kg. The lower canopy vigor (TGN = 0.83) and reduced fruit mass traits (FS = 1.97) explain the limited productivity, despite nutritional advantages. Overall result demonstrated that, Cluster 1 represents high-yielding genotypes combining superior canopy architecture, fruit morphology, and mineral content; Cluster 0 exhibits balanced canopy and quality traits suited for stable yield under semi-arid conditions; while Cluster 2 contributes fruit quality and nutritional diversity at the expense of yield. The parallel coordinate trends affirm that yield is strongly coupled with vegetative vigor and nutrient assimilation traits, while color and biochemical traits drive fruit quality differentiation. The visualization clearly delineates three functional ideotypes; high-yielding (Cluster 1), structurally stable (Cluster 0), and quality-focused (Cluster 2) which together provide complementary parental resources for integrated breeding strategies in semi-arid regions.

**Table 6:**
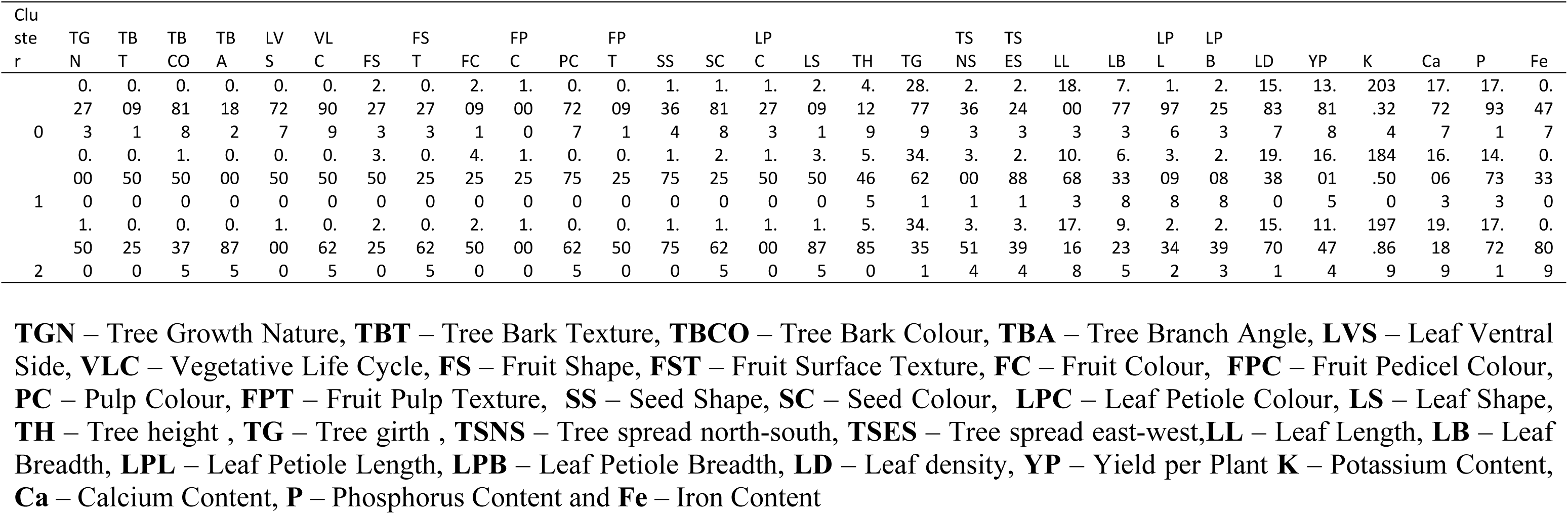
Cluster-wise mean distribution of morphological, fruit, seed, biochemical, and yield traits among *Annona reticulata* genotypes identified through multivariate clustering analysis.

**Figure 6:**
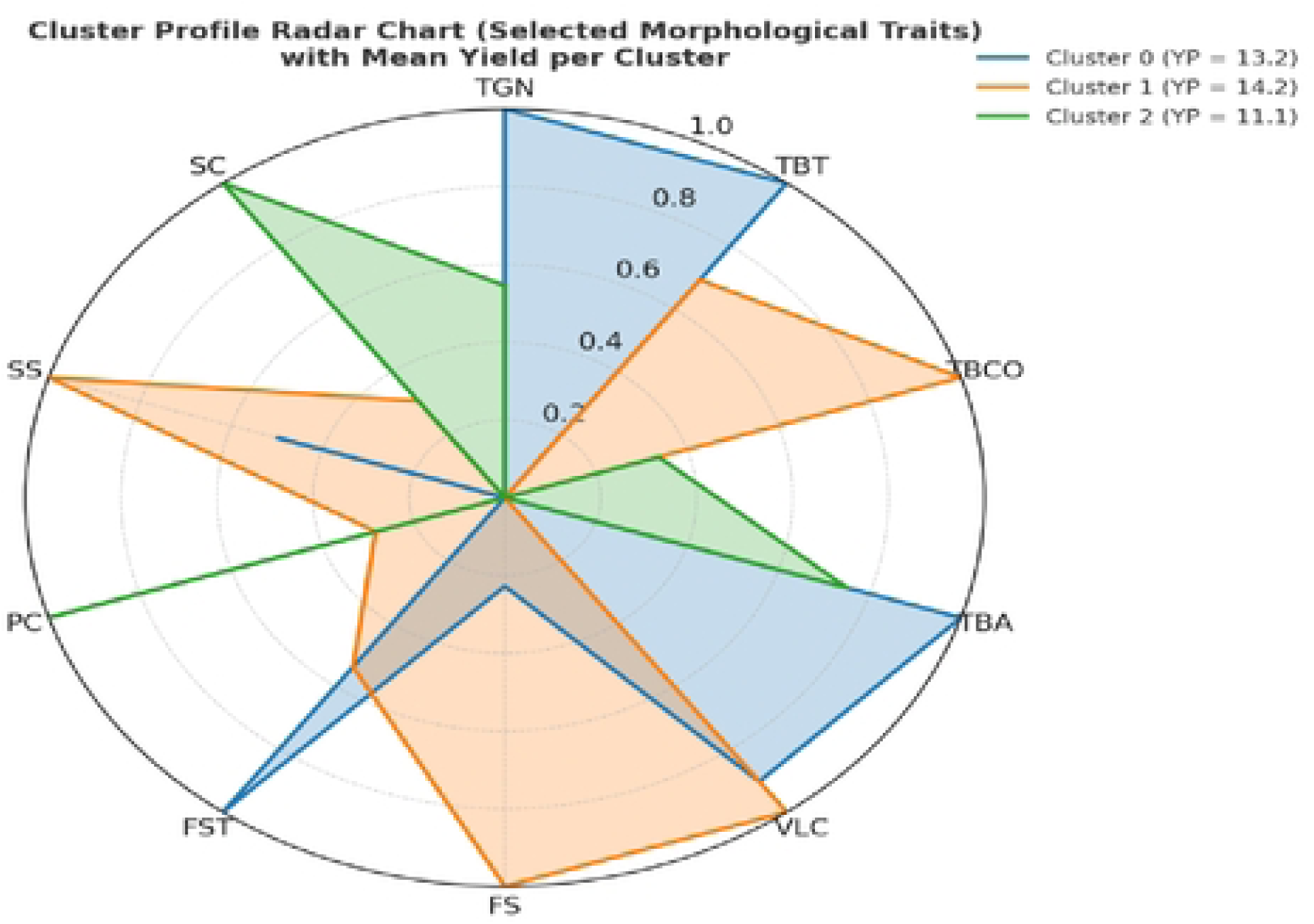
Cluster Profile Radar Chart showing normalized mean trait expression for three clusters of *Annona reticulate* genotypes based on morphological traits and mean yield (YP)

**Figure 7:**
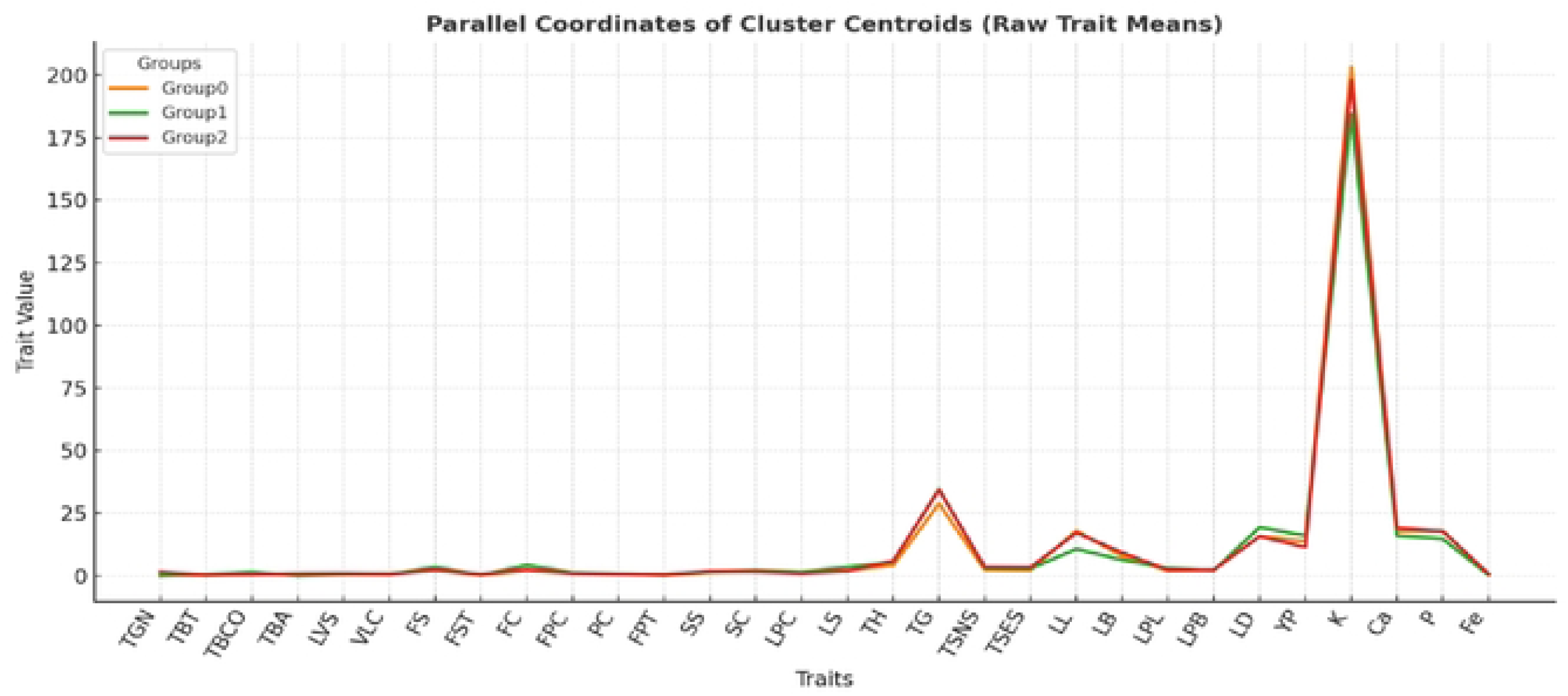
Parallel coordinates plot of cluster centroids showing raw mean values of morphological, fruit, seed, and biochemical traits across *Annona reticulata* genotype groups.

**Figure 8:**
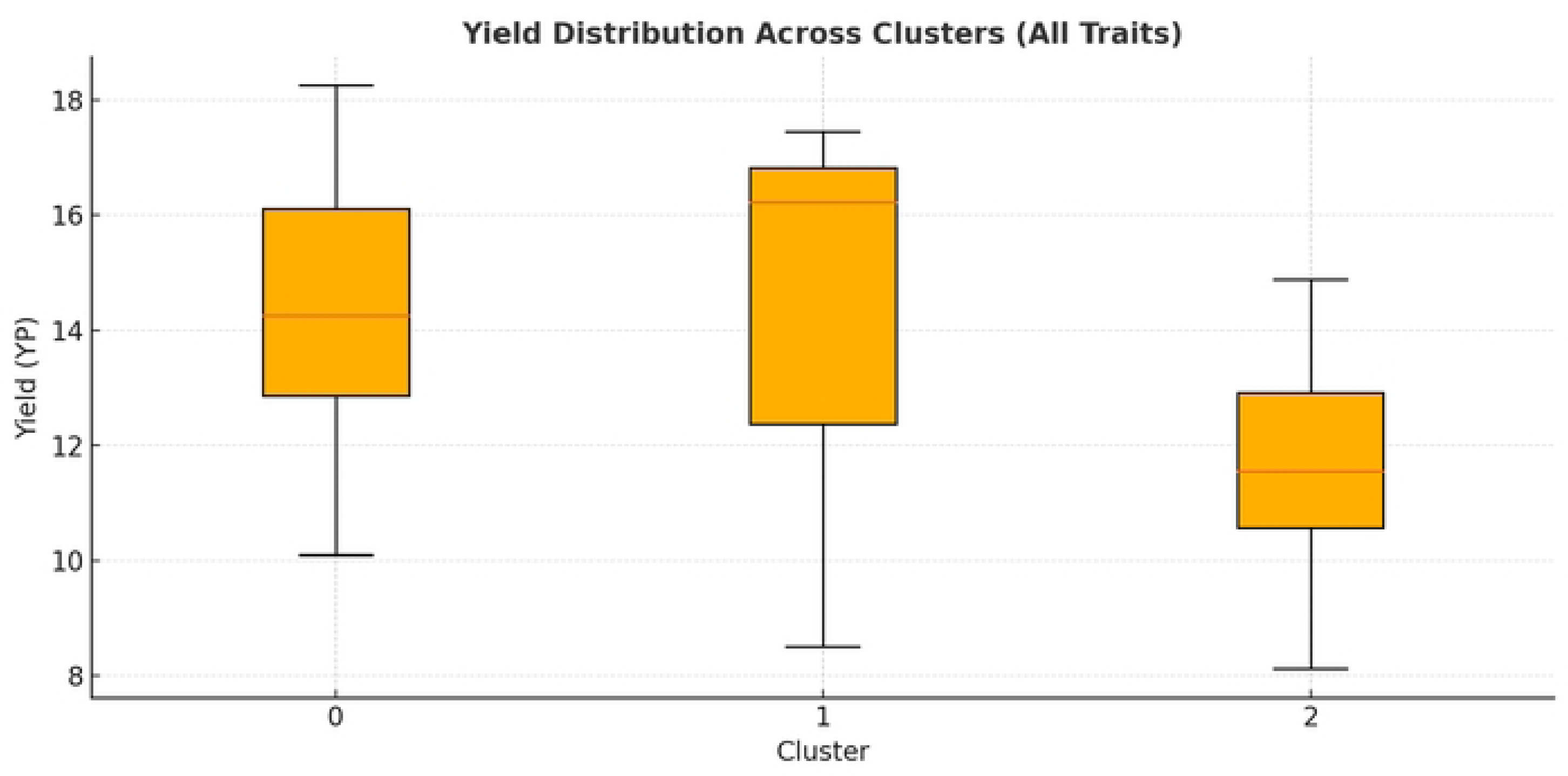
Boxplot illustrating yield per plant (YP) distribution among three multivariate-derived clusters of *Annona* reticulata genotypes. Cluster 1 exhibited the highest and most stable yield performance, with a median yield of 16.3 kg plant^-1^, followed by Cluster 0 (14.2 kg plant^-1^), while Cluster 2 recorded the lowest median yield (11.6 kg plant^-1^).

### The SHAP summary plot

The integration of SHAP-derived mean importance with trait variability (CV%) revealed a weak negative correlation (r = –0.25, p = 0.18), showing that moderately variable traits contribute most consistently to yield prediction in *Annona reticulata (*Fig. 9 and table7). LB, LL, LD, FS, and LPL displayed the highest SHAP importance, highlighting their dominant roles in assimilate partitioning and canopy efficiency. Importantly, these traits also showed moderate CV values, suggesting that they are both yield-responsive and genetically stable, making them reliable selection indices for breeding programs. Although, traits with high CV such as TBT, VLC, and TGN showed minimal SHAP impact, implying inconsistent or indirect influence on yield under semi-arid conditions. The combined SHAP–CV assessment therefore underscores that yield improvement is governed primarily by leaf and fruit morphological traits exhibiting intermediate variability, which balance predictive strength with phenotypic reliability.

**Figure 9:**
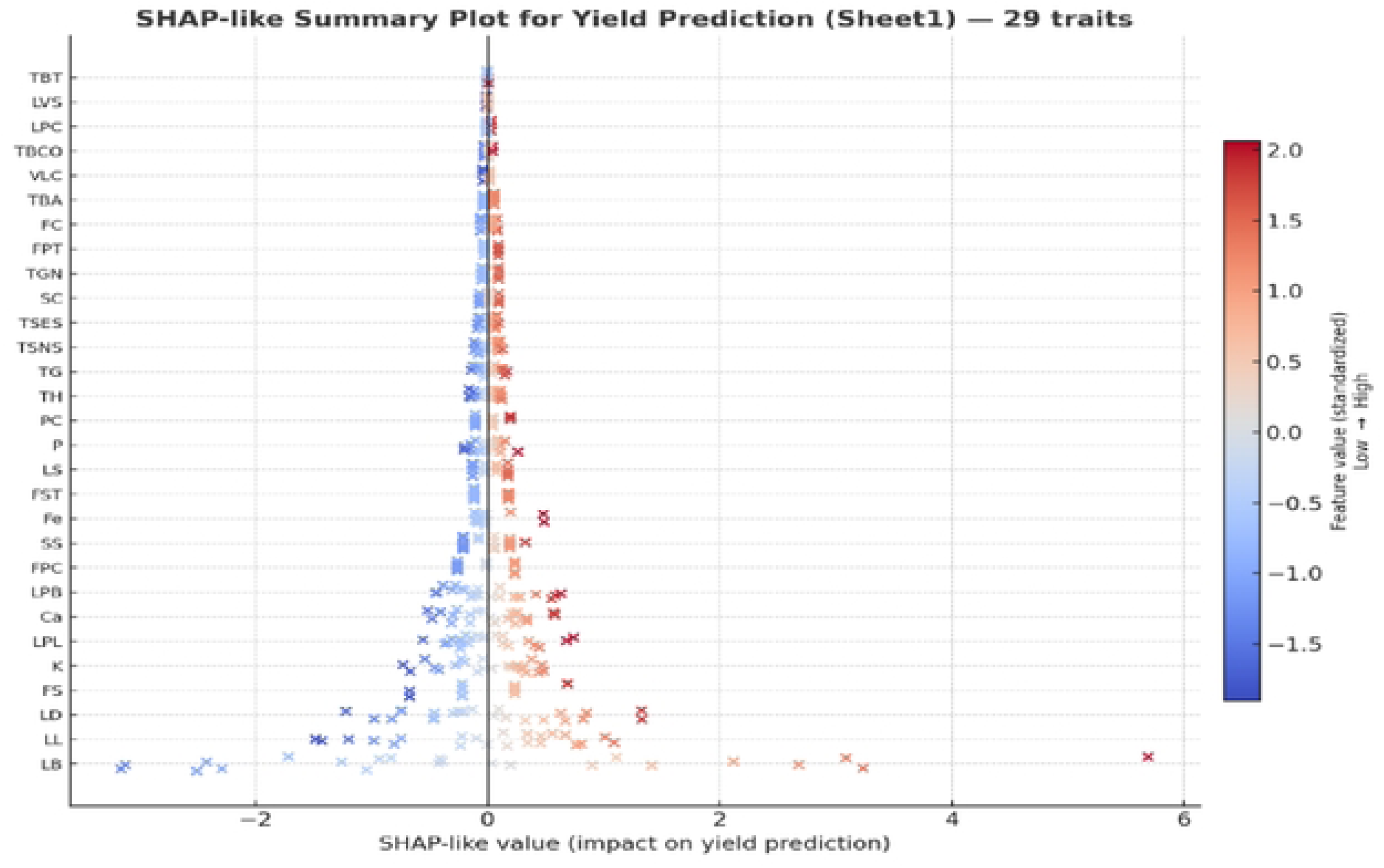
SHAP-based feature importance analysis for yield prediction using morphometric descriptors of *Annona reticulata* genotypes.

**Table 7.**
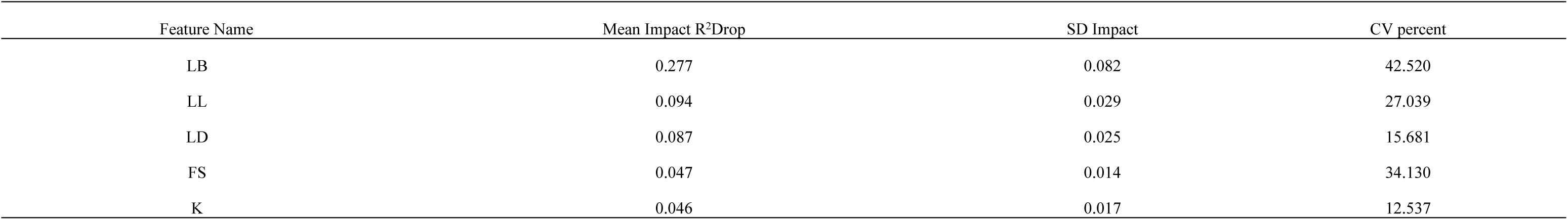
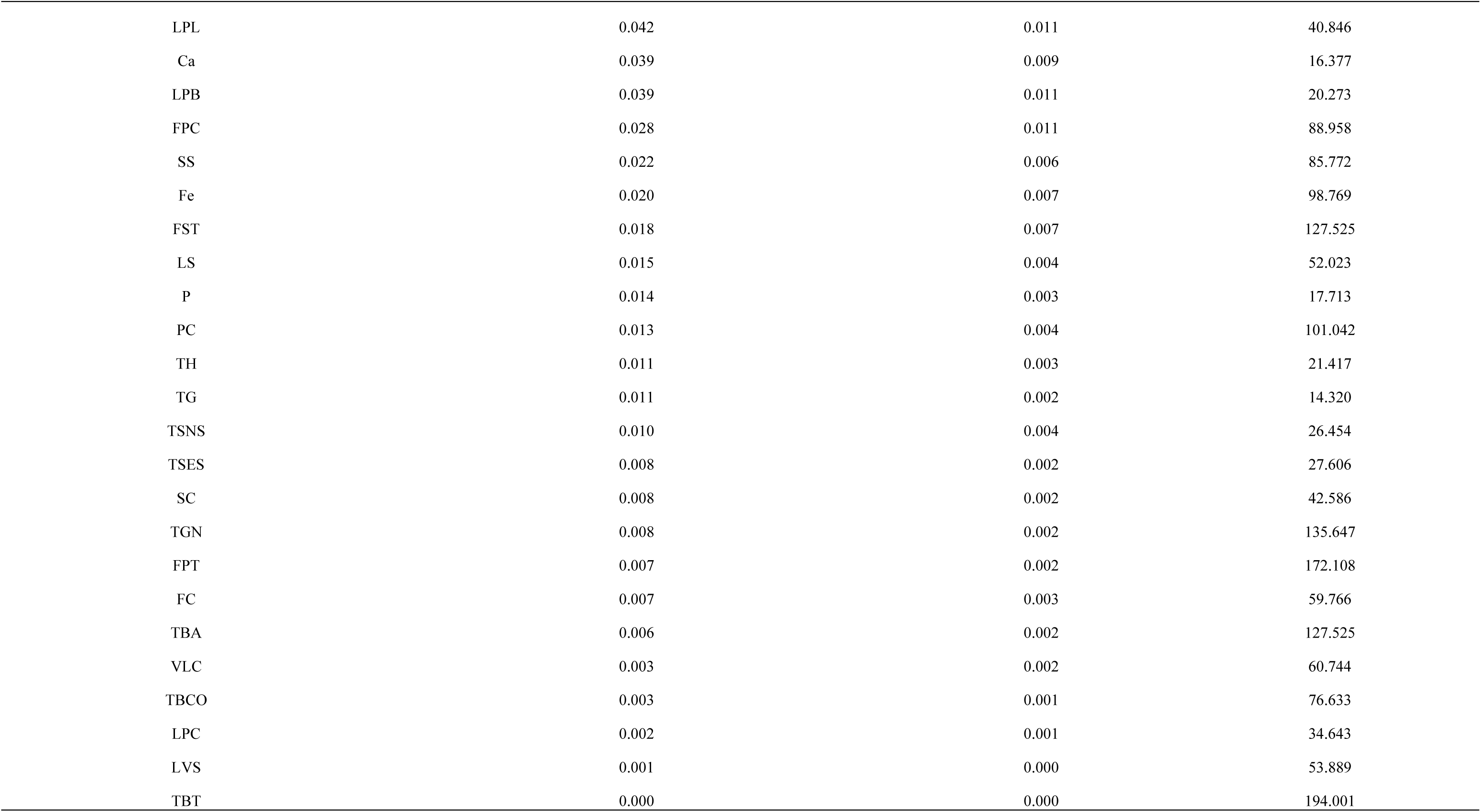
SHAP-derived feature importance and coefficient of variation (CV%) for identifying stable and yield-influential morphometric traits in *Annona reticulata* genotypes.

| Feature Name | Mean Impact R<sup>2</sup>Drop | SD Impact | CV percent |
| --- | --- | --- | --- |
| LB | 0.277 | 0.082 | 42.520 |
| LL | 0.094 | 0.029 | 27.039 |
| LD | 0.087 | 0.025 | 15.681 |
| FS | 0.047 | 0.014 | 34.130 |
| K | 0.046 | 0.017 | 12.537 |
| LPL | 0.042 | 0.011 | 40.846 |
| Ca | 0.039 | 0.009 | 16.377 |
| LPB | 0.039 | 0.011 | 20.273 |
| FPC | 0.028 | 0.011 | 88.958 |
| SS | 0.022 | 0.006 | 85.772 |
| Fe | 0.020 | 0.007 | 98.769 |
| FST | 0.018 | 0.007 | 127.525 |
| LS | 0.015 | 0.004 | 52.023 |
| P | 0.014 | 0.003 | 17.713 |
| PC | 0.013 | 0.004 | 101.042 |
| TH | 0.011 | 0.003 | 21.417 |
| TG | 0.011 | 0.002 | 14.320 |
| TSNS | 0.010 | 0.004 | 26.454 |
| TSES | 0.008 | 0.002 | 27.606 |
| SC | 0.008 | 0.002 | 42.586 |
| TGN | 0.008 | 0.002 | 135.647 |
| FPT | 0.007 | 0.002 | 172.108 |
| FC | 0.007 | 0.003 | 59.766 |
| TBA | 0.006 | 0.002 | 127.525 |
| VLC | 0.003 | 0.002 | 60.744 |
| TBCO | 0.003 | 0.001 | 76.633 |
| LPC | 0.002 | 0.001 | 34.643 |
| LVS | 0.001 | 0.000 | 53.889 |
| TBT | 0.000 | 0.000 | 194.001 |

### The Random Forest

Model-agnostic feature importance analysis using a Random Forest regression combined with permutation-based R² reduction (Figure 10, 11) revealed that a few morphometric traits predominantly drive yield variation in *Annona reticulata*. The top-ranking predictors were LB, LL, and LD, which together contributed over 60 % of the total predictive variance. These canopy-related traits directly influence photosynthetic efficiency, transpiration dynamics, and assimilate supply to developing fruits, thereby enhancing fruit biomass and yield potential. Among reproductive traits, FS, FW, and TH also exhibited significant contributions, indicating the central role of fruit morphology and sink strength in determining productivity. Secondary factors such as FPC and SS contributed moderately, possibly reflecting genotype-specific reproductive adaptations under semi-arid stress. Traits such as TGH, TBT, and VLC exhibited negligible permutation impact, suggesting that they exert indirect or minimal influence on yield. The low importance of these traits may also reflect their relatively low variability or limited physiological linkage with reproductive output. Overall, this analysis confirms that yield in *A. reticulata*is primarily governed by leaf-area and fruit-mass–associated parameters, which represent the functional balance between source and sink components.

**Figure 10:**
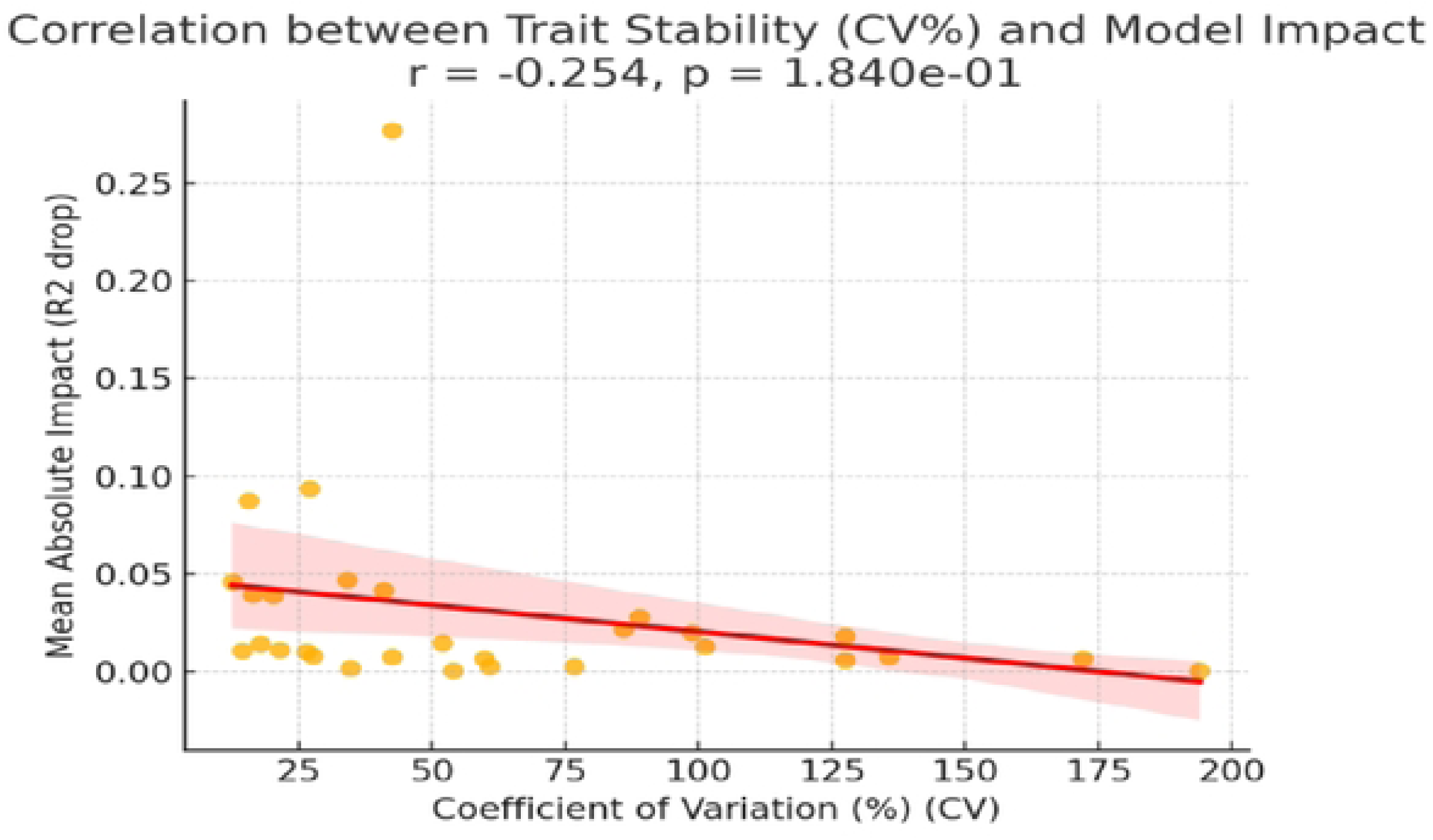
Correlation between mean absolute impact and trait stability (CV%) for yield-associated morphometric descriptors in *annona reticulate*.

**Figure 11:**
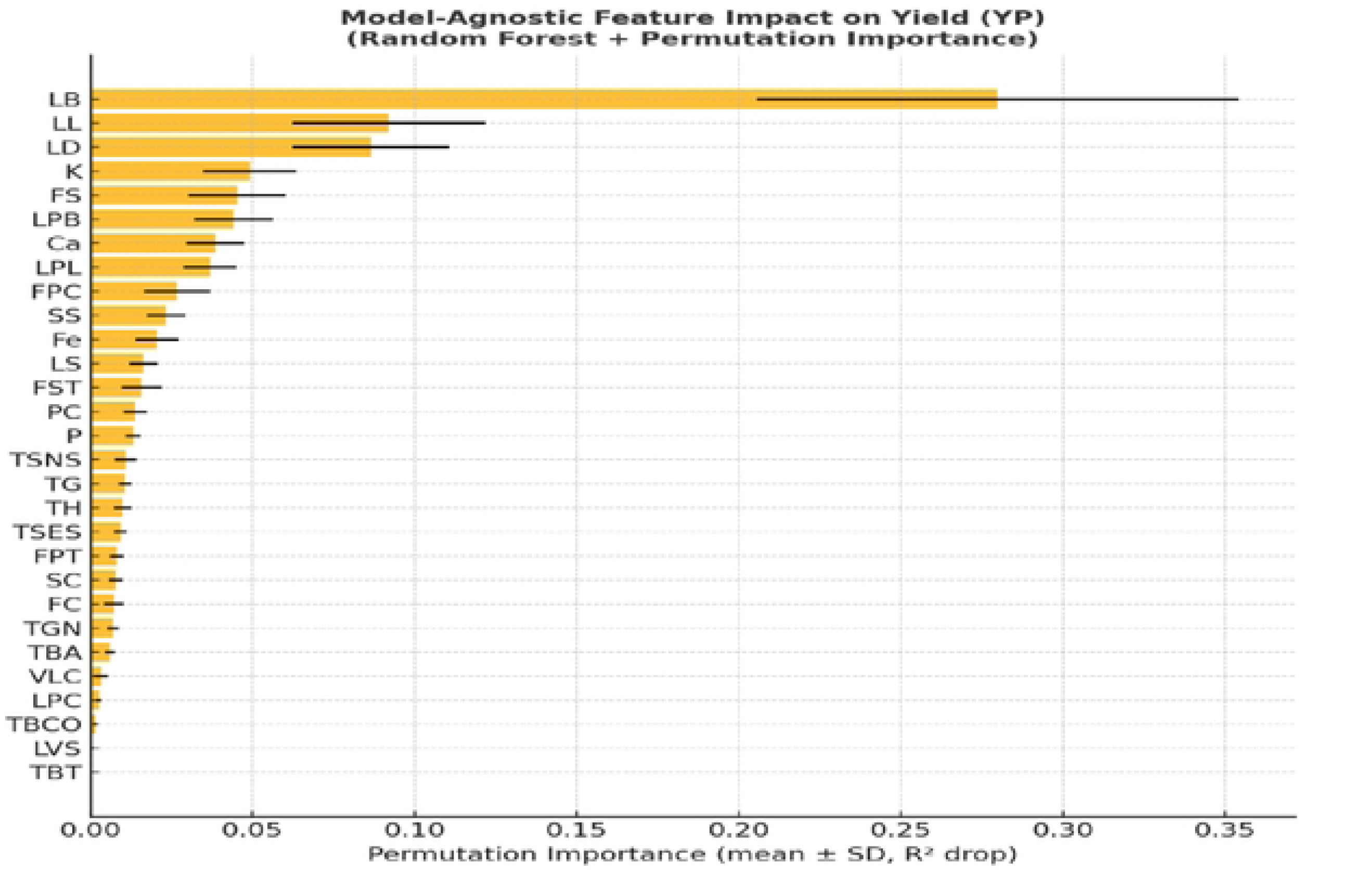
Model-agnostic feature importance analysis using Random Forest regression and permutation-based R^2^ reduction for yield prediction from morphological traits of 23 genotypes.

## Discussion

### Frequency distribution for morph-matrix characters

The observed morphological heterogeneity indicates a broad genetic base within the studied 23 ramphal genotype. Such variability is critical for crop improvement, as it enables the identification of ideotypes with balanced architecture, reproductive efficiency, and consumer-preferred fruit traits [**33,34**]. The morphological parameters align closely with the quantitative yield variation patterns, suggesting that morphological traits exert indirect but significant influence on fruit productivity (YP). Accessions exhibiting an upright to semi-spreading growth habit (74%) and smooth cylindrical bark texture (78%) demonstrated not only efficient canopy structure but also stable yield performance, as supported by the SHAP-like feature importance analysis where tree architecture traits contributed positively to yield prediction. The predominance of semi-deciduous genotypes (74%) ensures prolonged photosynthetic activity, optimizing assimilate partitioning toward fruit set and pulp development, a trend consistent with higher fruit weight, which were strongly associated with YP in the earlier model-based analysis [37]. Leaf traits also showed relationships with productivity. Genotypes with dark green glabrous leaves (78%) and lightly pubescent dorsal surfaces (100%) are likely to possess superior photosynthetic capacity and thermal tolerance, sustaining carbon assimilation during extended dry spells typical of semi-arid conditions. Such canopy efficiency corresponds to higher leaf length, breadth, and area, traits that exhibited strong positive contributions to yield in the SHAP analysis. The frequent occurrence of smooth fruit surfaces (61%) and creamy to pinkish pulp colour (87%) suggests a physiological balance between skin rigidity, and pulp hydration traits that favor assimilate translocation and fruit expansion, similar finding also reported by [16] in selection criterion for varietal development and commercialization.

Moreover, the predominance of spherical and oblong fruits (78%) with light brown to pinkish-yellow coloration indicates favorable sink strength and carotenoid metabolism, which have been linked with improved fruit biomass and yield stability. Accessions producing smooth pulp texture (74%) and dark to brown seed pigmentation (83%) reflect mature and well-filled fruits, further contributing to higher effective yield (YP). The integration of these traits with quantitative performance thus reveals that yield stability in *A. reticulata* is not governed by a single morphological factor, but rather by a synergistic expression of canopy architecture, foliar efficiency, and fruit surface pulp coordination and morphometric variability captured in structured datasets enables the machine learning algorithms to predict yield, fruit quality, and adaptability under different environments [8,38]. The integration of frequency distribution data with AI-driven feature-importance analyses provides a powerful framework for accelerating genotype screening, reducing experimental costs, and enabling data-driven breeding strategies [39–41]. The frequency distribution patterns not only underscore morphological richness present in ramphal but also establish the baseline phenotypic variability essential for deploying artificial intelligence in yield prediction and ideotype identification. By quantifying inherent diversity across vegetative and fruit traits, this approach strengthens the predictive capacity of ML models and supports the development of superior, climate-resilient genotypes. Consequently, the combined use of morphological distributions and AI analytics offers a robust pathway for advancing genetic improvement programs in underutilized perennial fruit crops.

### Descriptive statistics analysis

The descriptive statistical analysis and coefficients of variation (CV%) among morphometric and yield traits revealed substantial phenotypic diversity, indicating a broad genetic base within the *Annona reticulata* germplasm. The high CV observed in vegetative characteristics such as Tree Bark Texture, Tree Growth Nature, Tree Branch Angle, Fruit Surface Texture, Pulp Colour and Fruit Colour highlights the broad genetic variability within the studied 23 ramphal genotypes. Notably, fruit colour and texture is a key consumer-preference trait, and its substantial variation suggests ample scope for identifying superior market-oriented genotypes [5]. Such variability provides an important basis for selection, as traits with high diversity are generally more responsive to selection pressure in breeding programs. The observed variation also suggests strong genetic control with possible environmental modulation, offering considerable potential for the selection of ideotypes suited to specific orchard systems in semi-arid regions. In contrast, leaf morphometric traits (VLC, LDS, LVS, and LS) and elemental traits (Ca, P, and K) exhibited relatively lower coefficients of variation (CVs < 40%), indicating greater genetic stability and lower environmental sensitivity. Such stability is advantageous for maintaining physiological consistency across diverse growing environments [42]. Yield per plant (YP), marked by a relatively low CV (21.08%), which is consistent with the polygenic nature of yield as a complex trait, often buffered by compensatory mechanisms across vegetative and reproductive attributes. The relatively stable yield performance observed across genotypes highlights the potential of these germplasm for developing climate-resilient and high-yielding cultivars through selection and advanced modeling approaches, including machine learning-based yield prediction [43,44].

### Correlation analysis

Correlation analysis revealed that yield (YT) exhibited a positive correlation with LD (r = 0.39), followed by LPB (r = 0.36), TBCO (r = 0.30), FS (r = 0.27), VLC (r = 0.21), LS (r = 0.19), SS (r = 0.10), LPC (r = 0.10), and P (r = 0.19). These associations suggest that vigorous growth, regular bearing behaviour, and consumer-preferred nutrient-rich traits are important indicators of higher productivity. Similar patterns in other perennial crops confirm the central role of canopy architecture and reproductive efficiency in determining yield [45,46]. Importantly, fruit shape and phosphorous minerals functioned as a dual-purpose trait, linked to both consumer preference and productivity, making it a valuable predictive feature in AI-based modeling frame works. Although, weak to negative correlations of YT with morphological traits such as fruit surface texture, pulp colour, fruit colour, fruit pulp colour and seed colour highlight a trade-off between ornamental appeal and productivity. Such biological trade-offs, also noted in other horticultural crops, can be accounted for in AI-driven selection pipelines through the use of penalty functions or feature weighting, ensuring that yield optimization is not compromised by non-productive traits. Strong inter-correlations among vegetative descriptors, such as tree growth nature (TGN) with tree branch angle (TBA) and leaf length and breadth, reflect coordinated growth modules. These relationships, while not always directly linked to yield, can act as latent feature clusters that improve the explanatory power of machine learning models. These relationships indicate that excessive vegetative growth or traits linked to physiological stress and fruit quality variation may divert assimilates away from fruit development, thus; reducing yield. Collectively, the findings highlight that optimizing fruit structural traits while managing vegetative vigour could be crucial for improving yield potential. [47,16]. The strong negative correlation between TGN and TBCO (r = –0.66*) suggests contrasting lignin synthesis and pigment deposition patterns associated with upright tree forms, likely reflecting adaptive responses to photothermal or mechanical stress. Conversely, the positive association between LVS and Ca (r = 0.41*) indicates that leaf surface morphology and cuticular thickness may facilitate greater calcium accumulation, enhancing tissue rigidity and fruit firmness. The relationship between FST and PC (r = 0.42*) further demonstrates a structural–biochemical linkage in which surface traits align closely with pigment biosynthesis and cell-wall composition, influencing visual appeal and postharvest behavior. The moderate positive correlation between FC and YP (r = 0.30*) implies that enhanced pigment metabolism may be associated with improved assimilate partitioning and metabolic vigor. Additionally, the positive nutrient correlations among K, Ca, P, and Fe (r = 0.28–0.41*) highlight coordinated nutrient uptake and homeostasis, supporting both vegetative growth and reproductive output under marginal soil conditions. These finding would be helpful in future research into AI-driven predictive breeding pipelines to improve selection accuracy while balancing vegetative vigor and fruit quality. Exploring genotype-by-environment responses for nutrient-linked and structural traits may further enhance the development of resilient, high-yielding ideotypes

### Principal Component Analysis (PCA)

Principal Component Analysis (PCA) provided critical insights into the contribution of morphometric traits to yield variability in ramphal genotypes. The first two principal components (PC1 and PC2), which together explained 32.5% of the total variance, captured the dominant sources of phenotypic differentiation within the population, reflecting the multivariate complexity of trait–yield relationships. The dispersion of genotypes across the PCA biplot confirmed substantial morphological diversity, with clustering strongly influenced by TGN, TBA, and FST) on PC1, highlighting the significance of canopy configuration and external fruit morphology in defining genotypic variability. Genotypes positioned along the positive PC1 axis represented those with spreading canopies, wider branch angles, and smoother fruit surfaces, characteristics that favor efficient light interception and enhanced fruit set. These vegetative descriptors emerged as major contributors to variation, consistent with findings in other perennial fruit crops where canopy traits play a decisive role in productivity [16, 17]. Whereas, PC2 was primarily associated with SW, SL, SB and Yield per Plant (YP), signifying that this axis captured variation related to reproductive efficiency and sink strength. Genotypes with higher positive PC2 scores demonstrated elevated yield potential, reflecting optimal resource partitioning between vegetative and reproductive organs. The clustering of High-yielding genotypes (Top-1, Top-2, Top-3) within the positive quadrant of both PC1 and PC2 confirms that balanced morphological architecture and strong reproductive capacity jointly determine yield superiority. This highlights the importance of canopy architecture for maximizing light interception and photosynthetic efficiency. However, excessive vegetative vigour without reproductive balance can reduce fruiting, emphasizing the desirability of ideotypes with moderate vigour [15, 48].These findings are consistent with previous reports in tropical fruit crops, where multivariate components involving fruit size, surface texture, and pulp weight dominate phenotypic variance and predict yield potential [63].

### Clustering analyses

The clustering of ramphal genotypes classified them into three distinct groups, differing primarily in vegetative, reproductive, and fruit-related descriptors, which strongly influenced yield performance. Cluster 1 comprised genotypes with vigorous growth and balanced canopy traits, achieving a mean yield of 14.22 kg plant⁻¹, while tall, wide canopies enhanced light interception and assimilate production [17]. The wide yield range (10-14 kg/tree) indicated that excessive vigour without reproductive stability reduced consistency. Cluster 0 exhibited moderate yield performance (13.19 kg plant⁻¹) but demonstrated a balanced combination of structural stability and quality indicate robust canopy framework and well-developed reproductive structures, enhancing mechanical resilience and fruit set efficiency. These genotypes likely possess optimized photosynthate partitioning, ensuring stable productivity and consumer-preferred sweetness traits valuable for both table and processing markets [16]. The narrow yield variation within this group implied genetic uniformity but limited productivity potential. In contrast, Cluster 2, with the lowest mean yield (11.05 kg plant⁻¹), exhibited the highest Iron content (Fe = 3.26 mg 100 g⁻¹), signifying a nutrient-rich and visually superior nutrient rich fruit profile. The pronounced pigmentation and higher Fe concentration imply strong antioxidant potential and nutritional value, but smaller fruit explain the yield trade-off. Such genotypes, however, hold significance as donor parents for nutritional enhancement and biofortification initiatives. However, the adoption of efficient agrotechniques and improved orchard management practices may further enhance productivity [45]. In addition, compact trees facilitate high-density planting and reduce canopy-related stress, thereby improving orchard efficiency [47].

The convergence of PCA, SHAP-based feature importance, and cluster centroid analyses collectively demonstrates that canopy vigor, fruit morphology, and mineral assimilation traits are the principal determinants of yield in *Annona reticulata*. High-yielding genotypes consistently combined semi-spreading architecture, smooth fruit surface, and superior pulp recovery, supported by elevated Ca and P content, enhancing assimilate partitioning and fruit development. SHAP-based modeling further validated leaf size and fruit weight as top predictive features for yield, while PCA positioned these traits within the primary axes of variation. Collectively, these multivariate approaches confirm that morpho-physiological balance and nutrient efficiency underpin productivity and adaptability under semi-arid conditions.

### The yield distribution across clusters

The yield distribution analysis across clusters provides important insights into the relationship between morphological variability and productivity among ramphal genotypes. The marked differences among the clusters reinforce the grouping patterns identified by hierarchical clustering, demonstrating that canopy architecture and reproductive traits strongly influence yield performance [49]. Cluster 0, displaying the widest IQR (5.1 kg), encompasses genotypes with heterogeneous canopy forms and fruit characteristics, reflecting broader genetic variability and potential adaptability. The diversity within this cluster, as also indicated by PCA dispersion and intermediate centroid scores, highlights its value as a breeding pool for developing stress-resilient genotypes capable of maintaining productivity under fluctuating semi-arid conditions. Similar findings have been reported in other fruit crops where excessive vegetative growth compromises flowering and fruit set [48]. Cluster 1, narrowest interquartile range (3.7 kg), represents genotypes exhibiting strong canopy vigor, balanced branch architecture, and good fruit traits were positively loaded along PC1 and PC2 in the PCA. The reduced yield variability within this group demonstrates high genetic uniformity and efficient source sink dynamics, this outcome is consistent with trait-based analyses that highlighted limitations in bearing ability within these genotypes. Such stability, while beneficial for orchard uniformity, is less desirable from a commercial standpoint due to restricted yield potential. Cluster 2 clearly outperformed the other groups, combining both high yield levels and stability across replications. The narrow yield range and elevated median values indicate that genotypes within this cluster possess a favourable balance of compact canopy structure and superior reproductive efficiency. The limited presence of outliers underscores their robustness and adaptability. Cluster 2 clearly underperformed and showed a narrow IQR (3.0 kg), reflecting consistent though lower productivity among genotypes emphasizing quality-oriented traits such as high pulp colour intensity and mineral content (Fe and P). The parallel coordinate visualization confirmed that these accessions exhibit lower canopy and fruit size metrics but enhanced biochemical composition, reinforcing a trade-off between fruit quality and yield. The narrow yield range and elevated nutrient values indicate that genotypes within this cluster possess a favourable superior quality trait but small fruit size yet provide suitable line pool for breeding program for quality improvement. These results align with previous studies suggesting that ideotypes with optimized source-sink balance and efficient reproductive traits are better candidates for sustainable yield improvement [51,52]. The yield distribution pattern across clusters provides a quantitative validation of the phenotypic stratification observed in the PCA and cluster centroid analysis. The pronounced inter-cluster variation in median yield underscores that productivity in *Annona reticulata* is not randomly distributed but structured by specific combinations of morphological, physiological, and biochemical traits. The finding demonstrated that leaf breadth, leaf length, fruit weight, and pulp weight are the most influential predictors of yield. This multi-dimensional differentiation provides a clear framework for strategic parent selection in breeding programs targeting the integration of yield efficiency, structural stability, and nutritional enhancement in *ramphal* under semi-arid environments.

### The SHAP summary plot

The SHAP analysis offered valuable insights into the relative importance of morphological, phenological, and fruit-related traits in predicting yield of ramphal genotypes. The weak negative correlation between CV% and SHAP values (r = –0.25, p = 0.18) indicates that moderately variable traits exert more consistent effects on yield, highlighting the advantage of genetically stable traits. Leaf attributes especially LB, LL, and LD showed high SHAP impact with moderate CV%, confirming their role as dependable indicators of canopy efficiency and assimilate allocation. SHAP analyses further identified key leaf and fruit morphological traits (LB, LL, FS, LPL) as major positive contributors to yield, while traits like TGN, TBT, and VLC showed minimal influence, reflecting weaker direct roles in productivity. Traits like LB, LL, LB, LL, FS, LPL, and LD emerged as the most influential predictors, exerting strong positive effects on productivity. This suggests that balance canopy trees with favourable bearing patterns and desirable fruits attributes are consistently linked to higher yields, corroborating earlier machine learning-based studies in horticultural crops where canopy architecture and fruit quality jointly determined yield performance [50, 53]. Leaf related descriptors, particularly leaf length, leaf density also contributed substantially. These fine-scale leaf morphological traits likely serve as proxies for photosynthetic efficiency and assimilate partitioning, thereby ensuring yield stability, as previously reported in perennial fruit crops [54, 69,70]. Overall, SHAP-based interpretation confirmed that yield is shaped by a complex interplay of growth architecture, reproductive traits, fruit quality, and structural features. This highlights the value of explainable AI for prioritising breeding traits and identifying both beneficial attributes and unfavourable trade-offs in perennial horticultural species.

### The Random Forest

Model-agnostic feature importance derived from the Random Forest regression framework, validated through permutation-based R² reduction, demonstrated that a limited set of morphometric traits predominantly governs yield variability in *Annona reticulata*. The canopy traits leaf breadth (LB), leaf length (LL), and leaf diameter (LD) collectively explained over 60% of the total predictive variance, confirming their dominant role in determining yield and productivity. These traits directly modulate light interception, gas exchange, and carbon assimilation, forming the primary source components that sustain fruit growth. Their high predictive influence, combined with moderate coefficient of variation values, indicates that they are both yield-responsive and genetically stable, making them reliable selection indicators in predictive breeding pipelines [16,56, 67]. Reproductive attributes such as fruit shape (FS), pulp weight (PW), and pulp percentage (PP) further enhanced model performance, highlighting sink strength and assimilate utilization efficiency as critical determinants of yield potential as well as serving both as a biochemical indicator of quality and a proxy for maturity and consumer preference [58]. however, traits like tree growth habit (TGH), bark texture (TBT), and vegetative life cycle (VLC) exhibited non-significant permutation effects, suggesting this work as undependable and site-specific influences and endorsed that balanced canopy is more efficient in plant yield prediction and might be reflecting trade-offs between defense and yield [59, 68]. Seed and fruit stalk traits influenced postharvest quality but not yield directly. Collectively, yield per tree (YT) results from integrative interactions among canopy structure, reproductive efficiency, and physiological optimization. Integrating Random Forest with morphological evaluation enables identification of priority traits, offering a robust framework for ideotype selection and genetic improvement of wood apple [60, 65, 66].

Overall, the integrative multivariate and SHAP analyses demonstrated that yield variation in *genotype* is governed by the coordinated expression of morphological, physiological, and biochemical traits. High variability in Tree Growth Nature (TGN) and Branch Angle (TBA) revealed extensive architectural diversity, offering potential for canopy-based selection to improve light interception and fruit set. Stable expression of mineral traits (Ca, P, Fe) indicated heritable biochemical regulation contributing to fruit quality. PCA and clustering identified three ideotypes; Cluster 1 (high-yielding, semi-spreading, mineral-rich), Cluster 0 (structurally stable, adaptable), and Cluster 2 (nutrient-dense, quality-oriented) confirming genetic heterogeneity and provide wide genetic base for selection. SHAP analysis pointed out leaf breadth, fruit weight, and fruit weight as key yield predictors, emphasizing the synergy between canopy vigor and assimilate partitioning. Collectively, these findings establish that balanced vegetative vigor, efficient resource use, and mineral homeostasis underpin yield stability, offering a foundation for breeding high-yielding, nutritionally enhanced, and climate-resilient custard apple cultivars.

## Conclusion

This study provides a comprehensive integration of morphometric trait analysis and machine learning modelling for yield prediction in ramphal, an underutilized yet resilient fruit tree. The results revealed considerable morphological diversity across 23 genotypes, with key descriptors such as tree spread and growth, branch angle, fruit colour, leaf and seed attributes playing decisive roles in yield determination. Principal Component Analysis, Random Forest modelling, SHAP interpretation, and hierarchical clustering collectively highlighted that productivity is not governed by a single factor but rather by a balance of canopy architecture, reproductive efficiency, and fruit quality. Notably, compact trees with superior reproductive traits (Cluster 1) demonstrated both higher and more stable yields, underscoring their potential as ideotypes for breeding and orchard establishment. The convergence of traditional statistical approaches with explainable ML demonstrated the strength of combining multivariate datasets to uncover complex, non-linear relationships between traits and yield. Positive predictors such as leaf breadth, leaf length, leaf diameter (LD), tree spread and fruit colour, alongside negative contributors like tree growth habit, bark texture, and vegetative life cycle, provide clear trait-based guidelines for selection. These insights not only expand the scientific understanding of ramphal productivity but also establish a framework for precision breeding and climate-smart orchard management. Therefore, the study concludes that yield prediction through morphometric characters represents an essential step toward precision horticulture. Future research should emphasize integrating morphometric data with real-time climate monitoring, remote sensing, and hybrid machine learning models to enhance prediction accuracy and adaptability under changing climatic scenarios. This holistic approach will strengthen decision-making in fruit production systems, ensuring improved productivity, profitability, and resilience of fruit crops.

## Acknowledgments

Research was supported by the Indian Council of Agricultural Research, Department of Agricultural Research and Education, Government of India is dully acknowledged. We are thankful to the Director, ICAR-CIAH, Bikaner, Rajasthan, India for providing the necessary facilities to complete this study.

The study did not involve the collection of plant, animal, or other materials from a natural setting

## Summary

**Fig 12:**
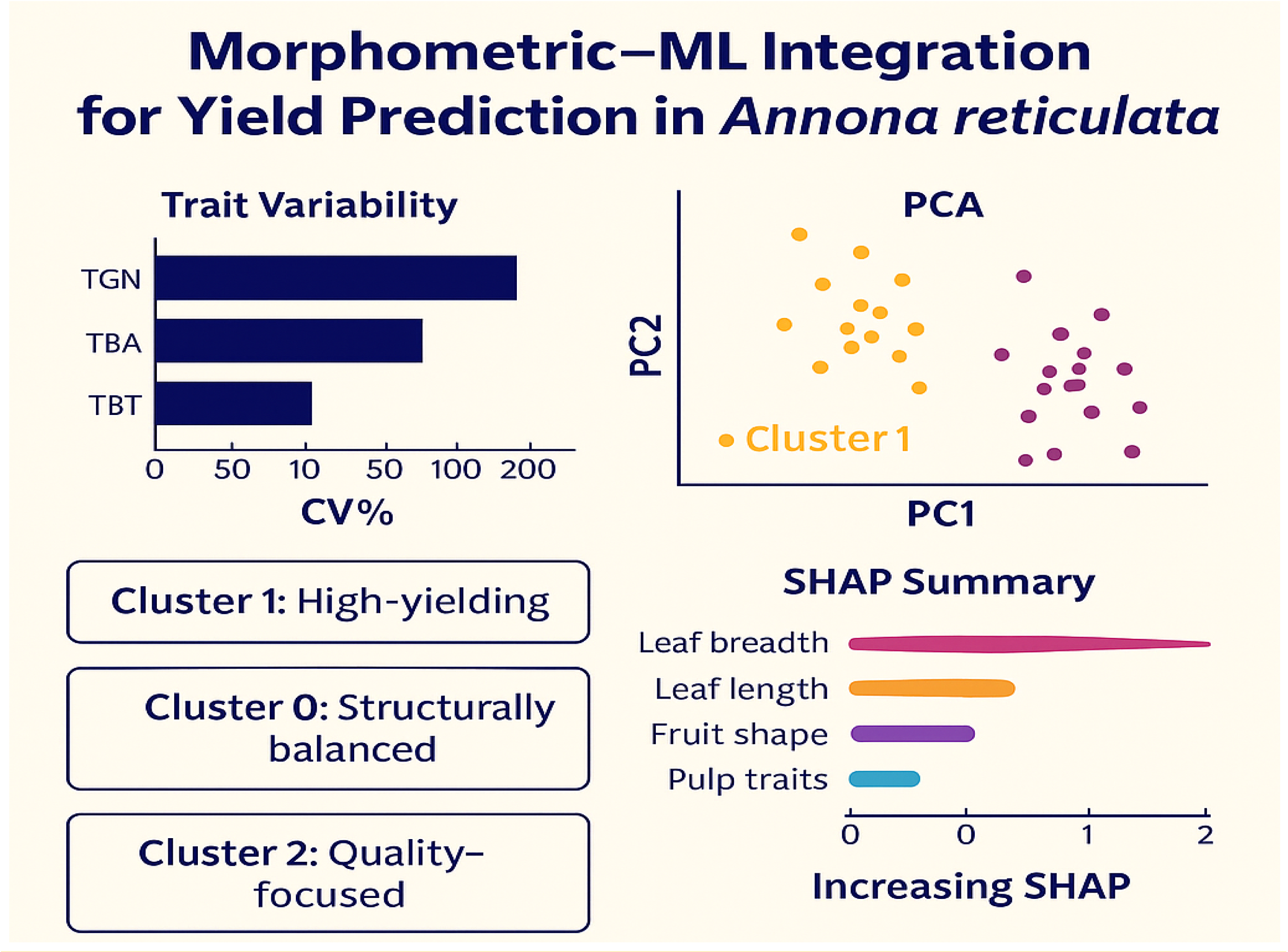
Integrated morphometric + ML framework enhances selection of high-yielder resilient genotypes

